# Immunometabolic analysis of primary murine Group 2 Innate Lymphoid Cells: a robust step-by-step approach

**DOI:** 10.1101/2024.12.16.628689

**Authors:** Sai Sakktee Krisna, Rebecca C. Deagle, Nailya Ismailova, Ademola Esomojumi, Audrey Roy-Dorval, Frederik Roth, Gabriel Berberi, Sonia V. del Rincon, Jörg H. Fritz

## Abstract

Group 2 Innate Lymphoid Cells (ILC2s) have recently been shown to exert key regulatory functions in innate and adaptive immune response networks that drive the establishment and progression of type 2 immunity and its associated pathologies. Although mainly tissue resident, ILC2s and their crosstalk within tissue microenvironments influences both local and systemic metabolism. In turn, the metabolic status shapes the diverse ILC2 phenotypes and effector functions. Hence, deciphering the metabolic networks of ILC2s is essential in understanding ILC2’s roles in health as well as pathophysiologies. Here we detail a framework of experimental approaches to study key immunometabolic states of primary murine ILC2s and link them to phenotypes and functionality. Utilizing flow cytometry, Single Cell ENergetIc metabolism by ProfilIng Translation inhibition (SCENITH) as well as the Seahorse platform we provide a framework that allows in-depth analysis of cellular bioenergetic states to determine the immunometabolic wiring of ILC2. Linking immunometabolic states and networks to ILC2 phenotypes and effector functions will allow in-depth studies that assess the potential of novel pharmaceutics to alter ILC2 functionality in experimental and clinical settings.

## 1. Introduction

Large extracellular helminths have been suggested to constitute a major evolutionary driving force of the effector mechanisms of type 2 immune responses, conferring protection from parasites invading barrier tissues (1, 2). However, the precise factors that shape the magnitude and quality of type 2 immune cell responses remain incompletely understood. Interestingly, chronic helminth infections have been shown to be associated with significant morbidity, including malnutrition, suggesting competition between parasites and host for metabolic resources, potentially leading to immunomodulatory consequences and alterations in protective type 2 immunity (3). Undeniably, an emerging body of evidence suggests that the immune system senses and utilizes nutrients and metabolites derived directly from the diet or produced by commensal or pathogenic microbes (4, 5). During the early phases of helminth infection, alarmin signals such as interleukin (IL)-25, IL-33, and thymic stromal lymphopoietin (TSLP) are released by non-hematopoietic cells in response to tissue damage and act to induce rapid proliferation and expansion of group 2 innate lymphoid cells (ILC2). Primarily found at mucosal barrier sites, ILC2 are transcriptionally and functionally poised innate type 2 effector immune cells that once activated can rapidly release large quantities of IL-5 and IL-13 to elicit eosinophilia, goblet cell hyperplasia, epithelial cell extrusion, and smooth muscle hypercontractility (6). Furthermore, it is gradually appreciated that ILC2 respond to changes in the richness and accessibility of microbially- as well as dietary-derived metabolites, such as vitamin A–derived retinoic acid (7), aryl-hydrocarbon receptor ligands (8), short chain fatty acids (9), succinate (10, 11), and iron (12), suggesting that ILC2 in addition to danger signals are poised to also respond to the broader metabolic milieu of barrier tissues (13). In addition, nutrients are essential components, providing fundamental metabolic substrates for the production the energy to fuel protein translation, cellular proliferation and immune cell effector functions (4, 5). Indeed, the ability of ILC2 to induce rapid effector functions depend on the ability to engage cell-intrinsic metabolic pathways to catabolize glucose, fatty and amino acids. Indeed, like T cells (14), ILC2s undergo metabolic reprogramming to meet the high energy demand imposed by cell proliferation and the activation of effector functions (15, 16).

In the resting state, ILC2 utilize branched chain amino acids (BCAAs) and arginine to support mitochondrial oxidative phosphorylation (OXPHOS) to sustain homeostatic functions (17). Following activation with IL-33, ILC2s become highly proliferative, relying on glycolysis and mammalian target of rapamycin (mTOR) to produce type 2 cytokines (18), while continuing to fuel OXPHOS with amino acids to maintain cellular fitness and proliferation (17, 19). Metabolic reprogramming during ILC2 effector differentiation also requires increased anabolic metabolism, leading to increased lipid droplet (LD) formation and requiring fatty acid oxidation (FAO) to fuel pathogenic allergic airway inflammation (20–23). In addition, the formation of lipid droplets is facilitated by glucose and was shown to be important to produce phospholipids that fuel ILC2 proliferation (21). Glucose was further shown to regulate the gene expression of diglyceride acyltransferase (DGAT1) and peroxisome proliferator-activated receptor gamma (PPARγ) through the mTOR pathway (21). While DGAT1 is driving LD formation (21), PPARγ promotes lipid uptake through CD36 expression (24–26). The increase in free fatty acid (FFA) uptake induced by ILC2 activation (20, 23) is fueled by group V phospholipase A2 (PLA2G5)-expressing macrophages (27). In addition, PLA2G5 intrinsically impacts IL-33 release from macrophages and the expression of the FFA receptor GPR40 on ILC2s (27). Interestingly, Atg5 deficiency lowered fatty acid metabolism gene expression induced by IL-33, and impaired FAO, type 2 cytokine production and cell fitness (22), suggesting that autophagy-mediated catabolic pathways are critical to sustain ILC2 metabolism.

In addition to sustaining sustain homeostatic functions (17), amino acid metabolism is essential for lung ILC2 activation and function. Arginine metabolism by arginase 1 (Arg1) is required to fuel aerobic glycolysis for ILC2 proliferation and cytokine production (28). ILC2 constitutively express high levels of multiple solute carriers, including Slc3a2, Slc7a5, and Slc7a8 (19), known to encode for large neutral amino acid transporter chains (29). Slc3a2 encodes the protein CD98 that heterodimerizes with other solute carriers to form active amino acid transporters, such as Slc7a5 and Slc7a8, which together form the surface amino acid transporters LAT1 and LAT2 (29). ILC2-intrinsic deletion of Slc7a5, and Slc7a8 impaired the proliferative and cytokine-producing capacity through tuning of mTOR signaling (19). In addition, catabolism of tryptophane (Trp) by the enzyme tryptophan hydroxylase 1 (Tph1) drives ILC2 effector functions (30). Moreover, methionine metabolism facilitates ILC2-driven inflammation through STAT3-dependent production of type 2 cytokines (31), underlining the various ways amino acid metabolism contributes to the regulation of ILC2 homeostasis, proliferation and cytokine production. Collectively, these insights highlight that fine-tuning metabolic pathways is critical for maintaining ILC2 homeostasis and regulating their effector functions, including cell fitness, proliferation and cytokine production. Here we detail a framework of experimental approaches to study key immunometabolic states of primary murine ILC2s and link them to phenotypes and functionality.

## Materials and methods

### 2.1 Mice

C57BL/6J mice were originally purchased from The Jackson Laboratory (Bar Harbor, Maine, USA) and bred in-house at McGill University under specific pathogen-free conditions with ad libitum access to food and water. Experiments were conducted with adult female mice (aged 8-12 weeks) in accordance with the guidelines and policies of McGill University and the Canadian Council on Animal Care.

### 2.2 Isolation of murine bone marrow and isolation of bone marrow-derived Group 2 Innate Lymphoid Cells

Adult female mice were anaesthetized with 5% isoflurane (Fresenius Kabi, Catalog No. CP0406V2) and euthanized via CO_2_ asphyxiation, followed by cervical dislocation as a confirmation of death. The hind legs were harvested, and the bulk of the muscles were removed with scissors to expose the leg bones. The femur and tibia bones were separated and cleaned with sterilized gauze (Fisher Scientific, Catalog No. 22-037-907) to remove any trace muscle or connective tissues, and briefly washed with 70% ethanol. Centrifugation was implemented to extract bone marrow from the femur and tibia bones. This centrifugation method began with the preparation of bone marrow extraction tubes, whereby a hole was punctured in the bottom of a 0.5 mL tube (Sarstedt, #72.737.002) with an 18-gauge needle (Becton Dickinson, Catalog No. 305196). These extraction tubes were sterilized by autoclave and each tube was set inside a sterile 1.5 mL collection tube (Progene, Catalog No. 87-B150-C) containing 100 µL of sterile Dulbecco’s Phosphate Buffered Saline (DPBS; Cytiva, Catalog No. SH30028.02). The bones were cut at the epiphysis on both ends to expose the bone marrow, and a single bone was vertically placed inside one 0.5 mL extraction tube. The 1.5 mL tubes containing the cut bones and extraction tubes were sealed and centrifuged at 1900*g for 5 minutes at room temperature. This allowed the bone marrow to be extracted through the punctured hole and into the collection tube with DPBS. After centrifugation the 0.5 mL tubes and hollow bones were discarded. The bone marrow was resuspended in the DPBS and transferred to a 50 mL tube (Fisher Scientific, Catalog No. 14-432-22), with a maximum of 20 bones (5 mice) per tube. Each 50 mL tube was filled to a volume of 50 mL with DPBS and centrifuged at 450*g for 5 minutes at 4°C to wash the bone marrow. The supernatant was discarded, and the bone marrow pellet was resuspended in 1 mL of Ammonium-Chloride-Potassium (ACK) lysis buffer (ThermoFisher Scientific, Catalog No. A1049201) for 20 to 30 seconds to remove red blood cells from the pellet. This reaction was neutralized by filling the tube to 50 mL with Fluorescence-Activated Cell Sorting (FACS) buffer made with 2% Fetal Bovine Serum (FBS; Wisent, Catalog No. 080150) in DPBS. The tube was centrifuged, and the pellet was washed twice with FACS buffer (450*g, 5 minutes, 4°C) to remove any trace of the lysis buffer. To prevent non-specific binding of antibodies to the fragment crystallization receptors (FcRs), the cells were blocked for 15 minutes on ice using a 1:10 dilution of in-house formulated “Fc-block” (supernatant of the 2.4G2 hybridoma producing the purified anti-mouse CD16/CD32 monoclonal antibody (mAb)) in FACS buffer. The bone marrow cells were stained with an antibody cocktail (Table 1) for 30 minutes on ice in the dark directly in the blocking solution. The stained bone marrow cells were washed twice in FACS buffer (450*g, 5 minutes, 4°C), filtered through a 70 µm pore strainer (Fisher Scientific, Catalog No. 22-363-548), and kept on ice prior to cell sorting. Bone marrow-derived ILC2 were sorted using a FACS Aria III and FACSAria Fusion (BD Biosciences) equipped with 405 nm, 488 nm, 561 nm (Fusion only) and 640 nm lasers using FACSDiva Version 6.0 (Aria III) or Version 8.0 (Fusion) based on the absence of lineage markers and the expression of ILC2 specific markers (Figure 1, Table 1).

**Figure 1.**
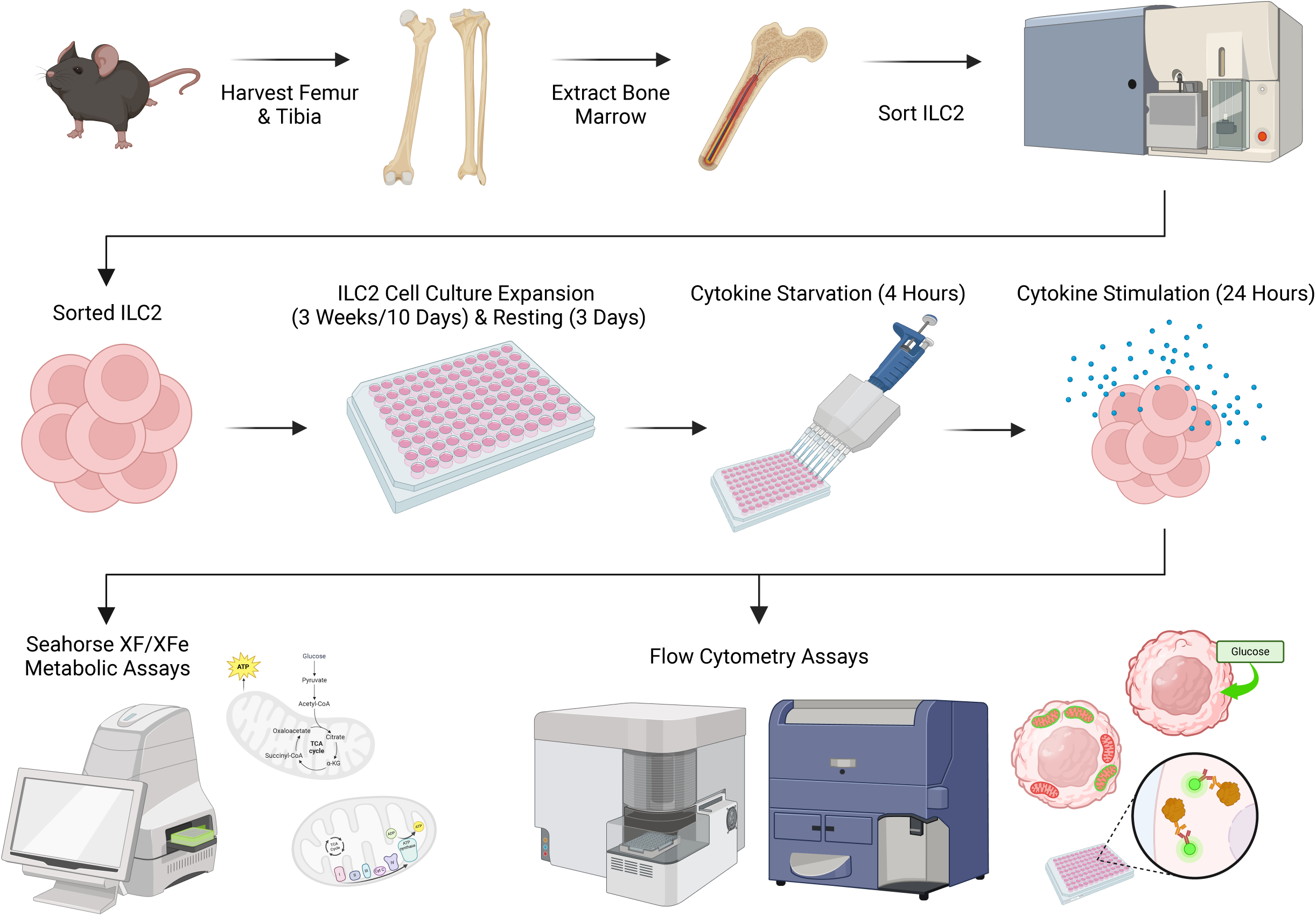
Schematic of experimental workflow. Femurs and tibias were harvested from female C57BL/6J wild-type mice and group 2 innate lymphoid cells (ILC2) were isolated from the bone marrow by flow cytometric cell sorting. ILC2 were subsequently cultured *in vitro* for expansion, followed by cytokine starvation (rest) and subsequent cytokine re-stimulation to determine metabolic activities by distinct flow cytometric assays as well as the Seahorse analyzer. Graph was created in BioRender. Krisna, S. (2024) BioRender.com/r16o748

**TABLE 1.** Group 2 Innate Lymphoid Cell (ILC2) sorting antibodies.

| Marker | Company | Fluorophore | Clone | Dilution | RRID or Catalogue Number |
| --- | --- | --- | --- | --- | --- |
| *Lineage Cocktail* |  |  |  |  |  |
| TCRα/β | eBioscience | PE | H57-597 | 1:200 | AB_466067 |
| TCRγ/δ | eBioscience | PE | eBioGL3 | 1:400 | AB_465934 |
| CD3e | eBioscience | PE | 145-2C11 | 1:400 | AB_465497 |
| Ly-6G/Ly-6C(Gr1) | eBioscience | PE | RB6-8C5 | 1:500 | AB_466046 |
| CD11b | eBioscience | PE | M1/70 | 1:500 | AB_2734870 |
| TER-119 | eBioscience | PE | TER-119 | 1:400 | AB_466043 |
| CD45R (B220) | eBioscience | PE | RA3-6B2 | 1:200 | AB_465672 |
| CD19 | eBioscience | PE | eBio1D3 | 1:200 | AB_657660 |
| NK1.1 | eBioscience | PE | PK136 | 1:50 | AB_466051 |
| CD5 | eBioscience | PE | 53-7.3 | 1:200 | AB_465524 |
| CD11c | eBioscience | PE | N418 | 1:100 | AB_465553 |
| FceR1 | eBioscience | PE | MAR-1 | 1:100 | AB_466029 |
| *ILC2+ Markers & Viability* |  |  |  |  |  |
| Ly-6A/E (Sca-1) | BioLegend | Alexa Fluor 488 | E13-161.7 | 1:200 | AB_756201 |
| CD25 | eBioscience | eFluor 450 | PC61.5 | 1:50 | AB_10671550 |
| CD117 (c-Kit) | eBioscience | APC | 2B8 | 1:25 | AB_469431 |
| Viability | eBioscience | 7-AAD | NA | 1:60 | 00-6993-50 |

### 2.3 Expansion and resting of murine bone marrow-derived Group 2 Innate Lymphoid Cells

After cell sorting, isolated ILC2 were cultured in 96-well round-bottom plates (VWR, Catalog No. CA62406-121) at 37°C in 5% CO_2_ with a seeding density of 2.5x10^4^ cells per well in 200 µL of complete cell culture media (Table 2) with cytokines for expansion, including: recombinant murine IL-2 (R&D Systems, Catalog No. 402-ML-100/CF), IL-7 (R&D Systems, Catalog No. 407-ML-200/CF), and IL-33 (R&D Systems, Catalog No. 3626-ML-010/CF) each at a concentration of 50 ng/mL and TSLP (R&D Systems, Catalog No. 555-TS-010/CF) at a concentration of 20 ng/mL. Every 2 days the cell culture was “split” whereby the cells were resuspended in the 200 µL volume, and 100 µL of the cell suspension was transferred to a new well. Each well was returned to a final volume of 200 µL by adding 100 µL of freshly prepared expansion cell culture media to facilitate proliferation. ILC2 were expanded for a maximum of three weeks, after which the cells were consolidated into 50 mL tubes and washed twice in RPMI 1640 (450*g, 5 minutes, 4°C) to remove any trace of expansion cytokines. The cells were counted and then seeded at a density of 2.5x10^4^ cells per well in 96-well round-bottom plates in 200 µL of complete cell culture media without β-mercaptoethanol (Table 2) and with 10 ng/mL each of IL-2 and IL-7. The cells incubated with these cytokines for three days at 37°C in 5% CO_2_ to “rest” the cells. This 3-day period of reduced activity was necessary after 3 weeks of energy-demanding proliferation to bring the cells down to a homeostatic baseline level of metabolic activity.

**TABLE 2.** Group 2 Innate Lymphoid Cell (ILC2) cell culture and experimental media recipes.

| Reagent | Company | Catalog Number | Final Concentration | Volume |
| --- | --- | --- | --- | --- |
| *Complete Cell Culture Media* |  |  |  |  |
| RPMI 1640 Medium | Cytiva | With Phenol Red<br>SH30096.02<br>Phenol Red-Free<br>SH30605.01 |  | 45 mL |
| Fetal Bovine Serum (FBS) | Wisent | 080150 | 10% | 5 mL |
| L-Glutamine (200 mM) | Cytiva | SH30034.01 | 2 mM | 500 µL |
| Penicillin-Streptomycin (10,000 µg/mL) | Cytiva | SV30010 | 100 µg/mL | 500 µL |
| Gentamicin (10 mg/mL) | Sigma-Aldrich | G1272 | 50 µg/mL | 120 µL |
| β-Mercaptoethanol (55 mM)<br>*Omitted for resting and experiments* | Gibco | 21985023 | 50 µM | 50 µL |
| *Agilent™ Seahorse XF Cell Mito Stress and XF Real-Time ATP Rate Assay* |  |  |  |  |
| Seahorse XF DMEM Medium | Agilent | 103575-100 |  | 48.5 mL |
| Seahorse XF Glucose (1M) | Agilent | 103577-100 | 10 mM | 500 µL |
| Seahorse XF Pyruvate (100 mM) | Agilent | 103578-100 | 1 mM | 500 µL |
| Seahorse XF Glutamine (200 mM) | Agilent | 103579-100 | 2 mM | 500 µL |
| *Agilent™ Seahorse XF Glycolysis Stress Assay* |  |  |  |  |
| Seahorse XF DMEM Medium | Agilent | 103575-100 |  | 49.5 mL |
| Seahorse XF Glucose (1 M) | Agilent | 103577-100 | 2 mM | 500 µL |

### 2.4 Cell seeding and cytokine stimulation for experimental assays

After resting, ILC2 were consolidated into 50 mL tubes and washed twice in RPMI 1640 (450*g, 5 minutes, 4°C) to remove any trace of resting cytokines. Cells were counted and seeded at a density of 1.0x10^5^ in 100 μL of phenol red-free complete cell culture media (Table 2) either in a 96-well round-bottom plate (Sections 2.5-7, 2.10) or a Seahorse XFe96 cell culture microplate (Agilent Technologies, Catalog No. 103794-100; Section 2.8). Each condition for flow cytometric assays (Sections 2.5-7, 2.10) were plated in triplicate and each condition for Seahorse assays (Section 2.8) were plated in quintuplicate. The ILC2 incubated for 4 hours at 37°C in 5% CO_2_ without cytokines to starve the cells of cytokines, bringing them to a latent stage of activity prior to cytokine stimulation (Figure 1). The cytokine stimulation conditions for all experiments were the following: IL-2, IL-7, IL-33, IL-2+IL-33, and IL-7+IL-33. Each cytokine stimulation was prepared as a 2X concentrated solution (20 ng/mL) in phenol red-free complete cell culture media (Table 2). After cytokine starvation, 100 μL of the appropriate cytokine stimulation was added to the corresponding well for a total volume of 200 μL and a 1X concentration of cytokines (10 ng/mL). The ILC2 underwent cytokine stimulation for a period of 24 hours at 37°C in 5% CO_2_ prior to all further experimental analysis.

### 2.5 Glucose uptake by flow cytometry

The glucose uptake in ILC2 was measured using the glucose analog 2-(N-(7-Nitrobenz-2-oxa- 1,3-diazol-4-yl)Amino)-2-Deoxyglucose (2-NBDG; ThermoFisher Scientific, Catalog No. N13195) which was stored at -20°C at a stock concentration of 10 mM (5 mg lyophilised powder in 1.46 mL dimethyl sulfoxide (DMSO)). Immediately prior to experiments, the 10 mM stock of 2-NBDG was diluted 1:100 in warm phenol red-free complete media (Table 2) to a 2X concentrated solution of 100 μM. After 24 hours of cytokine stimulation, the cells were washed twice in phenol red-free RPMI 1640 (450*g, 5 minutes, 4°C), resuspended in 100 μL of phenol red-free media, and transferred to a conical-bottom 96-well plate (Sarstedt, Catalog No. 82.1583.001). A volume of 100 μL of the 2X concentrated 2-NBDG was added to each well (excluding unstained and viability controls) for a final 1X concentration of 50 μM, and the plate incubated with the glucose analog for 30 minutes at 37°C in 5% CO_2_. After incubation, the cells were washed once with phenol red-free RPMI 1640 and once with FACS buffer (450*g, 5 minutes, 4°C). The supernatant was discarded, and the cells were resuspended in 195 μL of FACS buffer prior to acquisition on the flow cytometer. Immediately before the sample was acquired on the flow cytometer, 5 μL of viability dye (7-AAD; eBioscience, Catalog No. 00-6993-50) was added to each sample (excluding the unstained control wells) and transferred to a 5 mL FACS tube (Fisher Scientific, Catalog No. 352008). Each sample was acquired using a LSRFortessa Cell Analyzer (BD Biosciences) until 20,000 events were recorded. 2-NBDG (465/540 nm) was acquired using the 488 nm laser and 7-AAD (535/617 nm) with the 561 nm laser. After sample acquisition, the files were exported in .fcs format from FACSDiva and analysed using FlowJo software (BD, Version 10.10.0). Debris, doublet, and dead cells (7- AAD^+^) were excluded via flow cytometry gating. The 2-NBDG and the forward scatter were analysed on the x- and y-axes, respectively.

### 2.6 Analysis of PGC-1*α* expression levels by intracellular flow cytometry

#### Viability staining, fixation, and permeabilization

After 24 hours of cytokine stimulation, the cells were resuspended and transferred to a conical-bottom 96-well plate. The cells were washed twice with DPBS (450*g, 5 minutes, 4°C) and the supernatant was discarded. The fixable viability dye APC-eFluor780 (eBioscience, Catalog No. 65-0865) was stored at -80°C and diluted 1:1000 in DPBS. Excluding unstained controls, 50 μL of the diluted viability dye was added to each well and incubated on ice in the dark for 30 minutes. After incubation, the cells were washed twice with FACS buffer (450*g, 5 minutes, 4°C). The fixation/permeabilization solution was made during the centrifugation time, whereby the Fixation/Permeabilization Concentrate (eBioscience, Catalog No. 00-5123-43) was diluted 1:4 in Fixation/Permeabilization Dilutant (eBioscience, Catalog No. 00-5223-56). The supernatant from the cell culture plate was discarded, and 100 μL of the fixation/permeabilization solution was added to all wells and incubated on ice in the dark for 30 minutes. During incubation, the Permeabilization Buffer (10X; eBioscience, Catalog No. 00-8333-56) was diluted 1:10 in deionized water (dH_2_O). The fixed and permeabilized cells were washed twice with 100 μL of the 1X permeabilization buffer (600*g, 5 minutes, 4°C). The 2.4G2 hybridoma blocking solution was prepared during the centrifugation time as previously described (Section 2.2). The supernatant from the cell culture plate was discarded and 50 μL of blocking solution was added to all wells, followed by 15-minute incubation on ice in the dark.

#### Intracellular staining of PGC-1α

While incubating cells in the blocking solution, the antibody dilutions were made in 1X permeabilization buffer. The PGC-1α antibody (Proteintech, Catalog No. CL488-66369) was diluted 1:1000 from its stock concentration of 1000 μg/mL to 1 μg/mL and the IgG1 isotype control antibody (Proteintech, Catalog No. CL488-66360-1) was diluted 1:200 from its stock concentration of 200 μg/mL to 1 μg/mL. After incubation, the cell culture plate was centrifuged (600*g, 5 minutes, 4°C) and the supernatant was discarded. A volume of 100 μL of antibody dilution was added to the appropriate wells, excluding unstained controls, and incubated for 30 minutes on ice in the dark. The stained cells were washed twice with 1X permeabilization buffer (600*g, 5 minutes, 4°C) and the supernatant was discarded. The cells were resuspended in 200 μL of FACS buffer and transferred to a 5 mL FACS tube.

#### Flow cytometry

Each sample was acquired using an Aurora Spectral Flow Cytometer (Cytek Biosciences) until 20,000 events were recorded. The directly conjugated PGC-1α and isotype control (493/522 nm) were acquired using the 488 nm laser and APC-eFluor780 (756/785nm) with the 633 nm laser. After sample acquisition, the files were exported in .fcs format from FACSDiva (BD Biosciences) and analysed using FlowJo software (BD Biosciences, Version 10.10.0). Debris, doublet, and dead cells (APC-eFluor780^+^) were excluded via flow cytometry gating. The PGC-1α and the forward scatter were analysed on the x- and y-axes, respectively.

### 2.7 Analysis of mitochondrial mass and membrane potential by flow cytometry Fluorescent label preparation

Total mitochondrial mass was measured using Mitotracker^TM^ Deep Red FM (ThermoFisher Scientific, Catalog No. M22426), which was stored at -20°C at a stock concentration of 1 mM (50 μg lyophilised powder in 91.98 μL DMSO). Mitochondrial membrane potential was measured using Tetramethylrhodamine, Methyl Ester, Perchlorate (TMRM; ThermoFisher Scientific, Catalog No. T668) which was stored at -20°C at a stock concentration of 10 mM (25 mg lyophilised powder in 5 mL DMSO). Ready-to-use TMRM aliquots were prepared at an intermediate concentration of 100 μM by diluting the dye 1:100 in DMSO and were stored at -20°C until needed for experiments. Immediately prior to experiments, the 1 mM Mitotracker^TM^ Deep Red FM stock solution was diluted 1:10 in warm phenol red-free RPMI 1640 for an intermediate concentration of 100 μM. Both Mitotracker^TM^ Deep Red FM and TMRM were finally diluted together 1:1000 in warm phenol red-free RPMI 1640 to a 2X concentrated solution of 100 nM for experimental conditions. Each dye was also prepared separately for single-variable controls. In addition, cell viability was measured using 4′,6-diamidino-2-phenylindole (DAPI; ThermoFisher Scientific, Catalog No. 62248) where only dead cell nuclei would be labelled. The 1 mg lyophilized stock was reconstituted in 1 mL of dH_2_O to make a 1 mg/mL stock solution and was stored at 4°C. An intermediate concentration of 10 μg/mL was made by diluting DAPI 1:10 in FACS buffer immediately prior to sample acquisition.

#### Mitochondrial staining

After 24 hours of cytokine stimulation, the cells were washed twice in phenol red-free RPMI 1640 (450*g, 5 minutes, 4°C), resuspended in 100 μL of phenol red-free media, and transferred to a conical-bottom 96-well plate. The 2X concentrated Mitotracker^TM^ Deep Red FM and TMRM dilution was added to each well (excluding single colour and viability controls) at a volume of 100 μL for a final volume of 200 μL and a final concentration of 50 nM for each dye. The cells incubated with the dyes for 30 minutes at 37°C in 5% CO_2_. After incubation, the cells were washed once with phenol red-free media and once with FACS buffer (450*g, 5 minutes, 4°C). The supernatant was discarded, and the cells were resuspended in 198 μL of FACS buffer prior to acquisition on the flow cytometer.

#### Flow cytometry and data export

Immediately before the sample was acquired on the flow cytometer, 2 μL of the intermediate DAPI solution was added to each sample (excluding the unstained control wells) for a final concentration of 1 μg/mL and the cell suspension was transferred to a 5 mL FACS tube. Each sample was acquired using a LSRFortessa Cell Analyzer (BD Biosciences) until 20,000 events were recorded. Mitotracker^TM^ Deep Red FM (644/665 nm) was acquired using the 633 nm laser, TMRM (548/573 nm) with the 561 nm, and DAPI (350/470 nm) with the 405 nm laser. After sample acquisition, the files were exported in .fcs format from FACSDiva and analysed using FlowJo software. Debris, doublet, and dead cells (DAPI^+^) were excluded via flow cytometry gating. Mitotracker^TM^ Deep Red FM and TMRM were analysed on the x- and y-axes, respectively.

### 2.8 Agilent^TM^ Seahorse Extracellular Flux Assay preparation

#### Calibration and cell culture reagents

Concurrent with the Seahorse XFe96 microplate preparation (Section 2.4), the base of the Seahorse XFe96 sensor cartridge (Agilent Technologies, Catalog No. 103792-100) was loaded with 200 μL of Seahorse XF Calibrant (Agilent Technologies, Catalog No. 100840-000) and incubated in a CO_2_-free incubator at 37°C for 24 hours to hydrate the lid of the sensor cartridge. The day of the experiment, assay media for the Seahorse experiments were freshly prepared and warmed to 37°C for experiments (Table 2). The Seahorse XF Mito Stress Assay and Seahorse XF Real-Time ATP Rate Assay used the same assay media composition where Seahorse XF DMEM medium was supplemented with L-glutamine, glucose, and pyruvate (Table 2). The Seahorse XF Glycolysis Stress Assay used the Seahorse XF DMEM medium supplemented with only L-glutamine (Table 2).

#### Cell culture microplate

After 24 hours of cytokine stimulation, the cell culture microplate was removed from the incubator and centrifuged at 450*g for one minute at room temperature. The 24-hour old cell culture media was carefully removed from the plate to minimize cell disruption and replaced with 180 μL of the freshly prepared assay media specific to the Seahorse assay being conducted (Table 2). The cell culture microplate incubated for one hour in a CO_2_-free incubator at 37°C. Incubation of both the sensor cartridge and the cell culture microplate in CO_2_-free conditions permitted proper diffusion of CO_2_ from the cells, medium, and plate; ensuring that all were properly de-gassed prior to running the assays.

#### Stock solutions

During this one-hour incubation time, the relevant assay media was used to make stock solutions of the different compounds associated with the Seahorse XF Mito Stress Test Kit (Agilent Technologies, Catalog No. 103015-100), the Seahorse XF Real-Time ATP Rate Assay Kit (Agilent Technologies, Catalog No. 103592-100), or the Seahorse XF Glycolysis Stress Test Kit (Agilent Technologies, Catalog No. 103020-100). The Seahorse XF Mito Stress Test Kit compounds included oligomycin, carbonyl cyanide-4 phenylhydrazone (FCCP), and rotenone + antimycin A; the Seahorse XF Real-Time ATP Rate Assay Kit included oligomycin and rotenone + antimycin A; and the Seahorse XF Glycolysis Stress Test Kit included glucose, oligomycin, and 2-Deoxy-Glucose (2-DG; Table 3). Each compound was resuspended several times by pipetting with the corresponding assay medium and were vortexed for one minute to ensure a thorough reconstitution of the compound (Table 3).

**TABLE 3.** Stock solution calculations for each compound from the Agilent^TM^ Seahorse XF Cell Mito Stress Assay, Real-Time ATP Rate Assay, and Glycolysis Stress Assay Kits.

| Compound | Volume of Assay Medium | Stock Concentration |
| --- | --- | --- |
| *Agilent™ Seahorse XF Cell Mito Stress Assay* |  |  |
| Oligomycin | 630 µL | 100 µM |
| FCCP | 720 µL | 100 µM |
| Rotenone+Antimycin A | 540 µL | 50 µM |
| *Agilent™ Seahorse XF Real-Time ATP Rate Assay Kit* |  |  |
| Oligomycin | 420 µL | 150 µM |
| Rotenone+Antimycin A | 540 µL | 50 µM |
| *Agilent™ Seahorse XF Glycolysis Stress Assay Kit* |  |  |
| Glucose | 3000 µL | 100 mM |
| Oligomycin | 720 µL | 10 µM |
| 2-DG | 3000 µL | 500 mM |

#### Loading concentrations

The stock solutions were used to make a 10X concentrated solution of each compound with the corresponding assay media (Table 4). Each compound solution associated with the assay being performed was loaded into a specific port in the lid of the sensor cartridge (Port A, B, C, or D), which would be at a final 1X concentration after its timed injection during the assay (Table 4, Section 2.9). Any remaining compound solutions were discarded after the ports in the sensor cartridge lid were properly loaded. The Seahorse XFe96 sensor cartridge was returned to the CO_2_-free incubator until it was time to run the assay.

**TABLE 4.** Working concentration solutions and volumes of compounds for the Agilent^TM^ Seahorse XF Cell Mito Stress Assay, Real-Time ATP Rate Assay, and Glycolysis Stress Assay.

| Reagent | Stock Solution Volume | Assay Media Volume | Loading Concentration (10X) | Volume Added to Port | Final Concentration (1X) |
| --- | --- | --- | --- | --- | --- |
| *Agilent™ Seahorse XF Cell Mito Stress Assay* |  |  |  |  |  |
| Oligomycin | 450 µL | 2550 µL | 15 µM | 20 µL (Port A) | 1.5 µM |
| FCCP | 600 µL | 2400 µL | 20 µM | 22 µL (Port B) | 2.0 µM |
| Rotenone/<br>Antimycin A | 300 µL | 2700 µL | 5 µM | 25 µL (Port C) | 0.5 µM |
| *Agilent™ Seahorse XF Real-Time ATP Rate Assay* |  |  |  |  |  |
| Oligomycin | 450 µL | 2550 µL | 15 µM | 20 µL (Port A) | 1.5 µM |
| Rotenone/<br>Antimycin A | 300 µL | 2700 µL | 5 µM | 22 µL (Port B) | 0.5 µM |
| *Agilent™ Seahorse XF Glycolysis Stress Assay* |  |  |  |  |  |
| Glucose | 3000 µL | 0 µL | 100 mM | 20 µL (Port A) | 10 mM |
| Oligomycin | 300 µL | 2700 µL | 10 µM | 22 µL (Port B) | 1.0 µM |
| 2-DG | 3000 µL | 0 µL | 500 mM | 25 µL (Port C) | 50 mM |

### 2.9 Agilent^TM^ Seahorse Extracellular Flux Assay and data normalization

#### Parameter setting and calibration

The Seahorse XF/XFe Analyzer was turned on one hour prior to running the assays to stabilize. The Seahorse Wave Controller software (Agilent, Version 2.6) was opened on the computer, and the appropriate template file was selected (i.e.: Seahorse XF Cell Mito Stress Test, Glycolysis Stress Test, or Real-Time ATP Rate Assay). Under *Assay Navigation,* the option *Group Definitions* was selected and then *Groups* to create sample names. Sample names were determined based on the cytokine stimulations. A map of the conditions in the cell culture plate was generated under *Plate Map,* where wells labelled A1, A12, H1, and H12 were excluded to account for background signal. The sample names under *Group* were selected and applied to the generated plate map. A *Project Name* was created under *Run Assay* to identify the type and date of the assay, followed by *Start Run* to begin the calibration process. When the software prompted *Load Calibrant Utility Plate,* the sensor cartridge was removed from the CO_2_-free incubator and placed in the Seahorse XF/XFe Analyzer by selecting *Open Tray.* The option *I’m Ready* was selected to begin calibration; lasting approximately 20 minutes.

#### Seahorse data acquisition

When calibration was complete, the lid of the sensor cartridge was retained by the machine, and the base of the sensor cartridge was removed from the loading tray and replaced with the base of the cell culture microplate. The well labelled A1 in the cell culture microplate was aligned with the top-left corner of the loading tray and the option *Load Cell Plate* was selected to close the loading tray. The appropriate assay was chosen in the software to begin the assay. During the runtime of the assay, the Seahorse XF/XFe Analyzer automatically performed a timed injection of the relevant compound from the corresponding port in the lid of the sensor cartridge and into the cell culture plate below. The message *Unload Sensor Cartridge* signalled that the assay was complete, and the lid of the sensor cartridge was removed from the machine. *Eject* was selected to remove the cell culture microplate from the Seahorse XF/XFe Analyzer. The prompt *Assay Complete!* appeared on screen and the file was saved for viewing results after data normalization.

#### Real-Time ATP rate assay

To quantify absolute ATP production, we used the Agilent Seahorse XF ATP Real-Time rate assay, which measures and quantifies the rate of ATP production from the glycolytic and mitochondrial system simultaneously. While OXPHOS consumes O_2_ driving the oxygen consumption rate (OCR), both pathways contribute to the acidification of the assay medium. Conversion of glucose to lactate through glycolysis is accompanied by extrusion of one H^+^ per lactate. In addition, the TCA cycle that fuels ETC/OXPHOS produces CO_2_, which also results in acidification of the assay medium. The summary of these metabolic reactions is the primary driver of changes in extracellular acidification (ECAR). Seahorse XF technology measures the flux of both H^+^ production (ECAR) and O_2_ consumption (OCR) simultaneously. By obtaining these data under basal conditions and after serial addition of mitochondrial inhibitors (oligomycin and rotenone/antimycin A), total cellular ATP production rates and pathway-specific (mitochondrial vs glycolytic, mitoATP and glycoATP, respectively) production rates can be measured. The series of calculations used to transform the OCR and ECAR data to ATP production rates is as follows: glycolytic ATP production rate calculation: During conversion of one molecule of glucose to lactate in the glycolytic pathway, 2 molecules each of ATP, H^+^ and lactate are produced. Considering the stoichiometry of the glycolytic pathway, the rate of ATP production in the glycolytic pathway (glycoATP production rate) is equivalent to glycolytic Proton Efflux Rate (glycoPER) and can as such be calculated using the same approach: glycoATP production rate (pmol ATP/min) = glycoPER (pmol H^+^/min). Mitochondrial ATP production rate calculation: The rate of oxygen consumption that is coupled to ATP production during OXPHOS can be calculated as the OCR that is inhibited by addition of the ATP synthase inhibitor, oligomycin: OCR_ATP_ (pmol O_2_/min) = OCR (pmol O_2_/min) - OCR_Oligo_ (pmol O_2_/min). Transformation of OCR_ATP_ to the rate of mitochondrial ATP production consists of: multiplying by 2 to convert molecules of O_2_ to oxygen (O) atoms consumed, then multiplying the P/O ratio, the number of molecules of ADP phosphorylated to ATP per atom of O reduced by an electron pair flowing through the electron transfer chain. The Seahorse XF Real-Time ATP Rate Assay uses an average P/O value of 2.75 for these calculations that was validated to accurately represent the assay conditions for a broad panel of cells under different fuels availabilities With these considerations, the rate of mitochondrial ATP production is calculated as: mitoATP Production Rate (pmol ATP/min) = OCR_ATP_ (pmol O_2_/min) * 2 (pmol O/pmol O_2_) * P/O (pmol ATP/pmol O). Finally, the total cellular ATP Production Rate is the sum of the glycolytic and mitochondrial ATP production rates: ATP Production Rate (pmol ATP/min) = glycoATP Production Rate (pmol ATP/min) + mitoATP Production Rate (pmol ATP/min).

#### Cell viability data acquisition

The cell culture microplate was immediately used for cell viability quantification via flow cytometry to perform data normalization from the Seahorse assays. A volume of 5 μL of 7-AAD viability dye was added directly to each well, the cells were resuspended and transferred to a 5 mL FACS tube. Each sample was acquired using an Aurora Spectral Flow Cytometer (Cytek Biosciences) until 20,000 events were recorded. After sample acquisition, the files were exported in .fcs format from FACSDiva (BD Biosciences) and analysed using FlowJo software (BD Biosciences, Version 10.10.0). Debris, doublet, and dead cells (7-AAD^+^) were excluded via flow cytometry gating and the number of live, single cells were exported as an Excel file (Microsoft Corporation, Version 16.88) in .xls format.

#### Seahorse data normalization and export

The number of live, single cells were imported to a spreadsheet under *Normalize* in the Wave software. The Seahorse assay data was normalized according to the viability data by selecting *Apply.* To generate the data output from the assay, the *Results* tab was selected, followed by selecting the file of interest. In the *Functions* tab, *Export* was selected, and an Excel document was generated for each assay by selecting either Seahorse XF Cell Mito Stress Test Report Generator, Seahorse XF Glycolysis Stress Test Report Generator, or Seahorse XF Real-Time ATP Rate Assay Report Generator. Each Excel file was reviewed, and the three most consistent technical replicates (out of the five) were chosen from the dataset and transferred to GraphPad for graph configuration and statistical analyses (Section 2.11).

### 2.10 SCENITH analysis of murine bone marrow-derived Group 2 Innate Lymphoid Cells

Single Cell ENergetIc metabolism by profilIng Translation inhibition (SCENITH) analysis was performed with bone marrow-derived ILC2.

#### Reagent preparation

Glucose metabolism was inhibited with the glucose analog 2-Deoxy-D-Glucose (2-DG; Sigma Aldrich, Catalog No. D6134-25G) which was stored at -20°C at a stock concentration of 2 M (25 g crystalline powder in 76.14 mL dH_2_O). Mitochondrial ATP synthesis was inhibited with the antibiotic oligomycin (Sigma Aldrich, Catalog No. 75351- 5MG) which was stored at -20°C at a stock concentration of 1 mM (5 mg lyophilized powder in 31.6 mL dH_2_O). The antibiotic puromycin (Sigma Aldrich, Catalog No. P7255-25MG) was used as a proxy for measuring protein synthesis, which was stored at -20°C at a stock concentration of 50 mg/mL (25 mg lyophilised powder in 500 µL dH_2_O). Immediately prior to experiments, each of these reagents were prepared as an intermediate 4X concentrated solution in complete cell culture media (Table 2). 2-DG was prepared at a concentration of 400 mM (1:5 dilution), oligomycin at 4 µM (1:250 dilution), and puromycin at 40 µg/mL (1:1250 dilution).

#### Metabolic inhibition and sample preparation

After 24 hours of cytokine stimulation, the cells were resuspended and transferred to a conical-bottom 96-well plate. The plate was centrifuged (450*g, 5 minutes, 4°C) and 100 µL of media was removed from each well, excepting the control wells which only removed 50 µL. In turn, 50 µL of the 4X concentrated solutions of either 2-DG, oligomycin, or a combination of the two were added to the respective wells for a total volume of 150 µL and incubated for 30 minutes at 37°C in 5% CO_2_. After incubation, 50 µL of the 4X puromycin was added to each well (excluding the negative control wells) for a final volume of 200 µL and incubated for an additional 15 minutes at the same conditions described above. The final 1X concentrations of 2-DG, oligomycin, and puromycin in the 200 µL volume were 100 mM, 1 µM, and 10 µg/mL, respectively.

#### Antibody staining for puromycin (protein synthesis proxy) detection

The cells were washed twice with DPBS (450*g, 5 minutes, 4°C) and then stained for viability with the fixable APC-eFluor780 dye, as described in Section 2.6. As described in Section 2.6, the cells were incubated with the dye for 30 minutes on ice in the dark, followed by fixation, permeabilization, washes in 1X permeabilization buffer, and blocking in 2.4G hybridoma. The anti-puromycin antibody (Millipore-Sigma, Catalog No. MABE343-AF488) was diluted 1:1000 from its stock concentration of 0.5 mg/mL to 0.5 μg/mL. After incubation, the cell culture plate was centrifuged (600*g, 5 minutes, 4°C) and the supernatant was discarded. A volume of 100 μL of antibody dilution was added to the appropriate wells, excluding unstained controls, and incubated for 30 minutes on ice in the dark. The stained cells were washed twice with 1X permeabilization buffer (600*g, 5 minutes, 4°C) and the supernatant was discarded. The cells were resuspended in 200 μL of FACS buffer and transferred to a 5 mL FACS tube for flow cytometric acquisition.

#### Flow cytometry

Each sample was acquired using an Aurora Spectral Flow Cytometer (Cytek Biosciences) until 20,000 events were recorded. The directly conjugated anti-puromycin (493/522 nm) was acquired using the 488nm laser and APC-eFluor780 (756/785 nm) with the 633 nm laser. After sample acquisition, the files were exported in .fcs format from FACSDiva (BD Biosciences) and analysed using FlowJo software (BD Biosciences, Version 10.10.0). Debris, doublet, and dead cells (APC-eFluor 780^+^) were excluded via flow cytometry gating. Anti-puromycin and forward-scatter were analysed on the x- and y-axes, respectively.

### 2.11 Flow cytometric and statistical analysis

Flow cytometry analysis was performed using FlowJo software (BD, Version 10.10.0). Geometric mean fluorescence intensities (GeoMFI or gMFI) of the signals were calculated, triplicates were averaged per stimulatory condition and presented in bar graphs with all flow cytometry data represented as mean ± standard deviation (SD). Histograms and bar graphs were created using FlowJo and Prism softwares (Graphpad, Version 9) respectively. Statistical analysis was performed as ordinary one-way ANOVA and post-hoc Tukey’s multiple comparison tests to obtain statistical significance (*p*-values) between experimental conditions. *p*-values below 0.05 were defined as statistically significant (\**p*≤0.05, \*\**p*≤0.01, \*\*\**p*≤0.001, \*\*\*\**p*≤0.0001).

## 3. Results

The fine-tuning of metabolic pathways is critical for maintaining ILC2 homeostasis and the regulation of cell fitness and effector functions. We therefore aimed to establish a comprehensive set of rapidly applicable experimental approaches to study metabolic activities of ILC2 using flow cytometry based assays in combination with the Seahorse analyser (Figure 1). The applied methods detail the level of glucose update, the magnitude of energy production obtained by glycolysis and oxidative phosphorylation, as well as the plasticity of ILC2 metabolic programming of these respective pathways. Furthermore, we addressed numerous facets of mitochondrial involvement in ILC2 metabolism including mitochondrial biogenesis, the total mass of mitochondria per cell, the respiration of mitochondria, and the performance of that respiration.

### 3.1 Determining glucose uptake by ILC2

To study metabolic activities of ILC2, we obtained primary murine sort purified bone marrow-derived ILC2 (Supplemental Figure 1A) that were further expanded *in vitro* as previously described (32, 33). IL-33 has been established as a key driver of ILC2 activation (34). IL-7 and IL-2 secreted by tissue-resident non-hematopoietic stromal cells as well as innate and adaptive immune cells, respectively, have been shown to act synergistically with IL-33 to induce proliferation and cytokine production (32, 35, 36). However, the effects of these cytokines (IL- 2, IL-7, IL-33) alone or in synergy (IL-2+IL-33, IL-7+IL-33) as they pertain to glucose metabolism in ILC2 remain incompletely understood.

We first aimed to establish glucose uptake by ILC2 at defined steady and activation states using 2-(*N*-(7-Nitrobenz-2-oxa-1,3-diazol-4-yl)Amino)-2-Deoxyglucose (2-NBDG), a fluorescent glucose analogue (Figure 2). To investigate how activating cytokines impact glucose uptake, bone marrow-derived ILC2 were incubated with IL-7, IL-2, IL-33 alone, or in combinations of IL-7+IL-33 or IL-2+IL-33. After 24 hours of cytokine stimulation, 2-NBDG was added to the cells, incubated for 30 minutes, and subsequently analysed by flow cytometry. Bone marrow-derived ILC2 treated with IL-7 (Figure 2A) or IL-2 alone (Figure 2B) exhibited moderate levels of exogenous glucose uptake, which considerably increased when treated in combination with IL-33. ILC2 treated with IL-33 alone displayed a higher uptake of exogenous glucose compared to IL-7 alone (Figure 2A), but not in regards to IL-2 alone (Figure 2B). In fact, the combination of IL-2+IL-33 exhibited the highest uptake of glucose compared to IL- 33 or IL-2 alone (Figure 2B). Collectively, these observations demonstrate that ILC2 are actively acquiring exogenous glucose at steady state but were found to take up the most glucose when synergistically activated by IL-2+IL-33.

**Figure 2.**
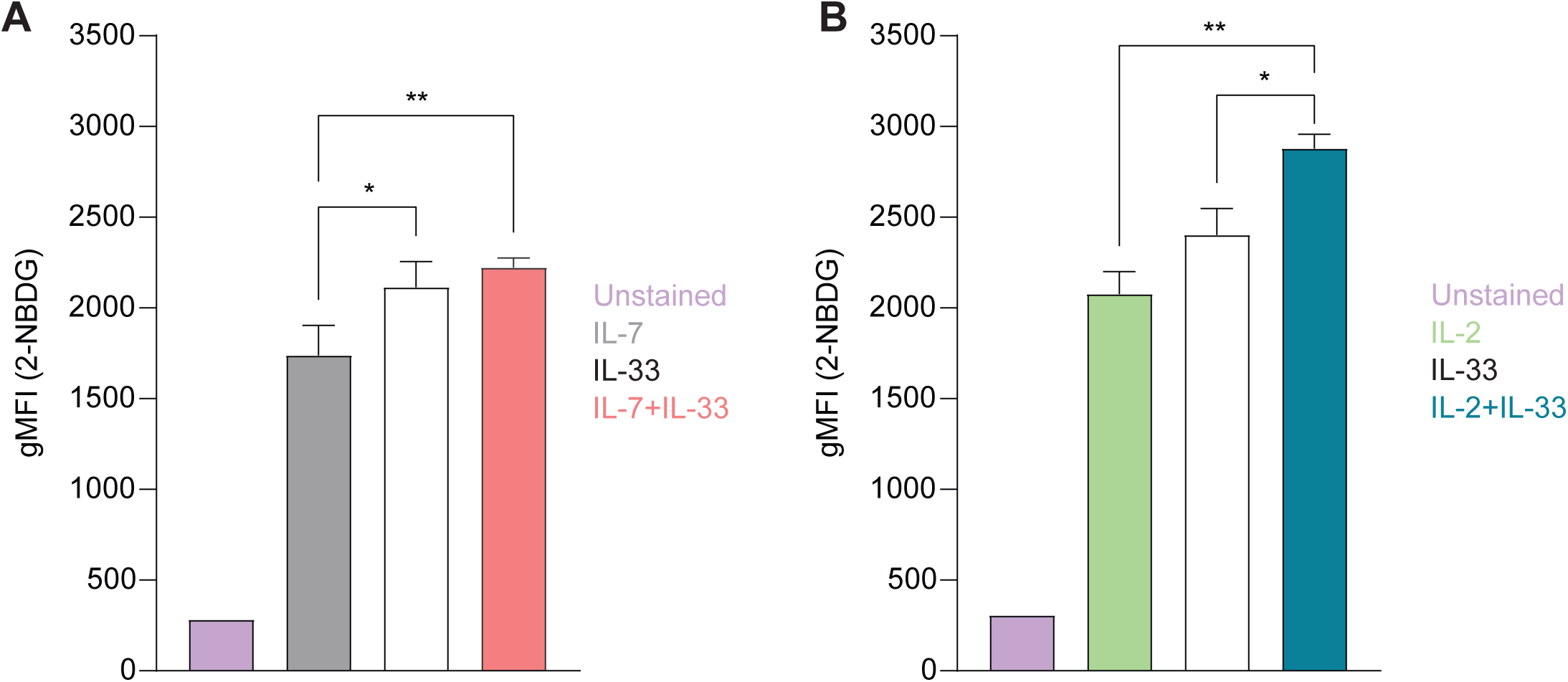
Group 2 innate lymphoid cells (ILC2) increase glucose uptake upon stimulation with activating cytokines. Bone marrow-derived group 2 innate lymphoid cells (ILC2) were stimulated with either IL-7 only, IL-33 only, or a combination of IL-7 and IL-33 **(A),** or IL-2 only, IL-33 only, or a combination of IL-2 and IL-33 **(B)**. All cytokines were applied at 10 ng/mL. 2-NBDG was added after 24 hours of cytokine stimulation and incubated for 30 minutes to assess capacity of glucose uptake by flow cytometric analysis, determining the geometric Mean Fluorescence Intensity (gMFI). ILC2 that were not incubated with 2-NBDG served as negative control (Unstained). Data reporting the treatment with IL-33 are the same for **(A)** and **(B)**. Data are shown as average ± standard deviation (SD) and are representative of three independent experiments. Statistical analysis was performed using one-way ANOVA followed by Tukey’s multiple comparisons test (p < 0.05 = *, p < 0.01 = **).

### 3.2 Quantification of mitochondrial biogenesis

Mitochondria are biosynthetic and bioenergetic organelles that also act as critical signalling platforms instructing decisions about cell proliferation, death, and differentiation. Mitochondria not only sustain immune cell phenotypes and functions, but to further fulfil appropriate metabolic demands can also switch from primarily being catabolic organelles generating ATP to anabolic organelles that in addition to ATP produce critical building blocks for macromolecule synthesis (37). Immune cells typically exhibit quiescent levels of metabolic activity at steady state and can shift to being highly metabolically active during the activation phase (5). This high energy demand triggers mitochondrial biogenesis to stimulate the production of more mitochondria and to replace mitochondria damaged by oxidative stress (38). The co-activator peroxisome proliferator-activated receptor gamma coactivator 1 alpha (PGC-1α) is characterized as a master regulator of mitochondrial biogenesis and oxidative metabolic pathways at both the transcriptional and post-translational levels (39). To investigate how cytokines known to drive ILC2 effector functions influence mitochondrial biogenesis, we first evaluated the expression levels of PGC-1α using intracellular flow cytometry. Bone marrow-derived ILC2 were incubated with IL-7, IL-2, IL-33 alone, or in combinations of IL- 7+IL-33 or IL-2+IL-33. After 24 hours of cytokine stimulation, the cells were fixed, permeabilized, and labelled using a directly conjugated antibody for PGC-1α followed by flow cytometric analysis. ILC2 treated with IL-7 (Figure 3A) or IL-2 alone (Figure 3B) exhibited moderate levels of PGC-1α expression, which considerably increased in the presence of IL-33. There was no significant elevation in PGC-1α expression between IL-33 alone and the combination of either IL-7+IL-33 (Figure 3A) or IL-2+IL-33 (Figure 3B). However, these synergistic combinations of activating cytokines displayed higher levels of PGC-1α expression compared to IL-7 (Figure 3A) or IL-2 alone (Figure 3B). These data suggest that the presence of IL-33 primarily influences PGC-1α expression to drive mitochondrial biogenesis rather than relying on the synergistic activation of cytokines known to influence ILC2 effector functions.

**Figure 3.**
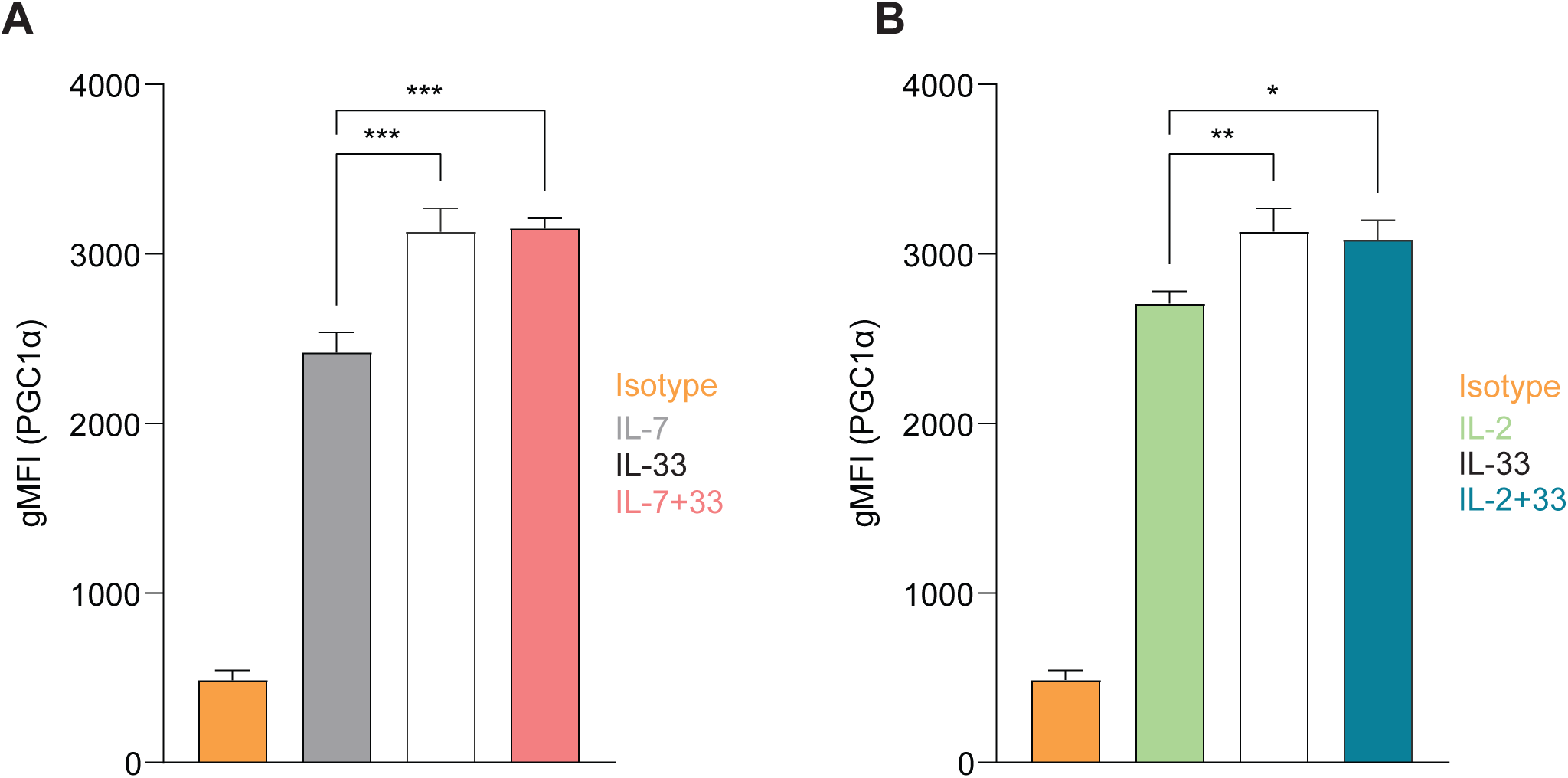
Group 2 innate lymphoid cells (ILC2) elevate expression of PGC-1a upon stimulation with activating cytokines. Bone marrow-derived group 2 innate lymphoid cells (ILC2) were stimulated with either IL-7 only, IL-33 only, or a combination of IL-7 and IL-33 **(A),** or IL-2 only, IL-33 only, or a combination of IL-2 and IL-33 **(B)**. All cytokines were applied at 10 ng/mL. After 24 hours of cytokine stimulation cells were harvested and the protein expression levels of the peroxisome proliferator–activated receptor gamma coactivator-1 alpha (PGC-1a) were assessed by intracellular flow cytometric analysis, determining the geometric Mean Fluorescence Intensity (gMFI). Isotype antibody stainings served as control (Isotype). Data reporting the treatment with IL-33 are the same for **(A)** and **(B)**. Data are shown as average ± standard deviation (SD) and are representative of three independent experiments. Statistical analysis was performed using one-way ANOVA followed by Tukey’s multiple comparisons test (p < 0.05 = *, p < 0.01 = **, p < 0.001 = ***).

### 3.3 Quantification of mitochondrial mass and active mitochondria

We next aimed to investigate mitochondrial metrics directly through the use of fluorescent functional dyes to be analyzed by flow cytometry. With the development of dyes such as MitoTracker^TM^ Deep Red, it is possible to label all mitochondria within living cells and quantify the total mass produced by ILC2 at defined steady and activation states. MitoTracker^TM^ Deep Red is a lipophilic carbocyanine-based dye that permeates the cell membrane to covalently bind thiol-reactive chloromethyl groups within the mitochondrial membrane, permitting all mitochondria to be fluorescently labelled (40). When glucose is acquired exogenously by the cell, the glucose is utilized by the mitochondria either through glycolysis or oxidative phosphorylation to produce energy in the form of adenosine triphosphate (ATP). While anaerobic glycolysis generates two ATP per glucose molecule, the aerobic process of oxidative phosphorylation produces 36 molecules of ATP (41). In brief, oxidative phosphorylation utilizes the electron transport chain (ETC) to drive protons (H^+^) against their concentration gradient out of the inner mitochondrial membrane space (41). This accumulation of H^+^ in the intermembrane space can then flow back through the ATP-generating component of the ETC, completing the energy production cycle (42). This difference in H^+^ concentration effectively creates both a pH and electrical gradient to generate a membrane potential in the mitochondria (41). This membrane potential, or polarization, can be exploited to label actively respirating mitochondria with other functional dyes, such as tetramethylrhodamine methyl (TMRM). TMRM is similar to MitoTracker^TM^ Deep Red in that they are both lipophilic cationic dyes so they will both be drawn into mitochondria across this charged gradient, however, TMRM exhibits a low binding affinity to mitochondrial proteins and functional groups (42). Effectively, MitoTracker^TM^ Deep Red will label all mitochondria that are present, whereas TMRM will preferentially label actively respirating mitochondria, and neither will interact with damaged mitochondrial membranes where this gradient is impaired.

To further analyze whether mitochondrial biogenesis was replacing potentially damaged mitochondria or increasing the overall mass during activation states, we optimized a flow cytometry-based protocol to rapidly quantify the total mass of mitochondria in live bone marrow-derived ILC2. To investigate how activating cytokines impact the overall mass of mitochondria synthesized by ILC2, cells were incubated with IL-7, IL-2, IL-33 alone, or in combinations of IL-7+IL-33 or IL-2+IL-33. After 24 hours of cytokine stimulation, the mitochondrial dye called MitoTracker^TM^ Deep Red was added to the cells, incubated for 30 minutes, and ILC2s were subsequently analysed by flow cytometry (Figure 4A). ILC2 treated with IL-7 (Figure 4B) or IL-2 alone (Figure 4E) exhibited moderate levels of mitochondrial mass, which considerably increased in the presence of IL-33 alone. While there was no significant elevation in mitochondrial mass between IL-7 alone and the synergistic combination of IL-7+IL-33 (Figure 4B), there was a slight increase between IL-2 alone and IL-2+IL-33 (Figure 4E). Interestingly, the mitochondria mass decreased between IL-33 alone and IL-7+IL- 33 (Figure 4B). These results suggest, similarly to the PGC-1α expression levels, that IL-33 influences the mitochondrial mass produced by ILC2 more than the synergistic effects of activating cytokines.

**Figure 4.**
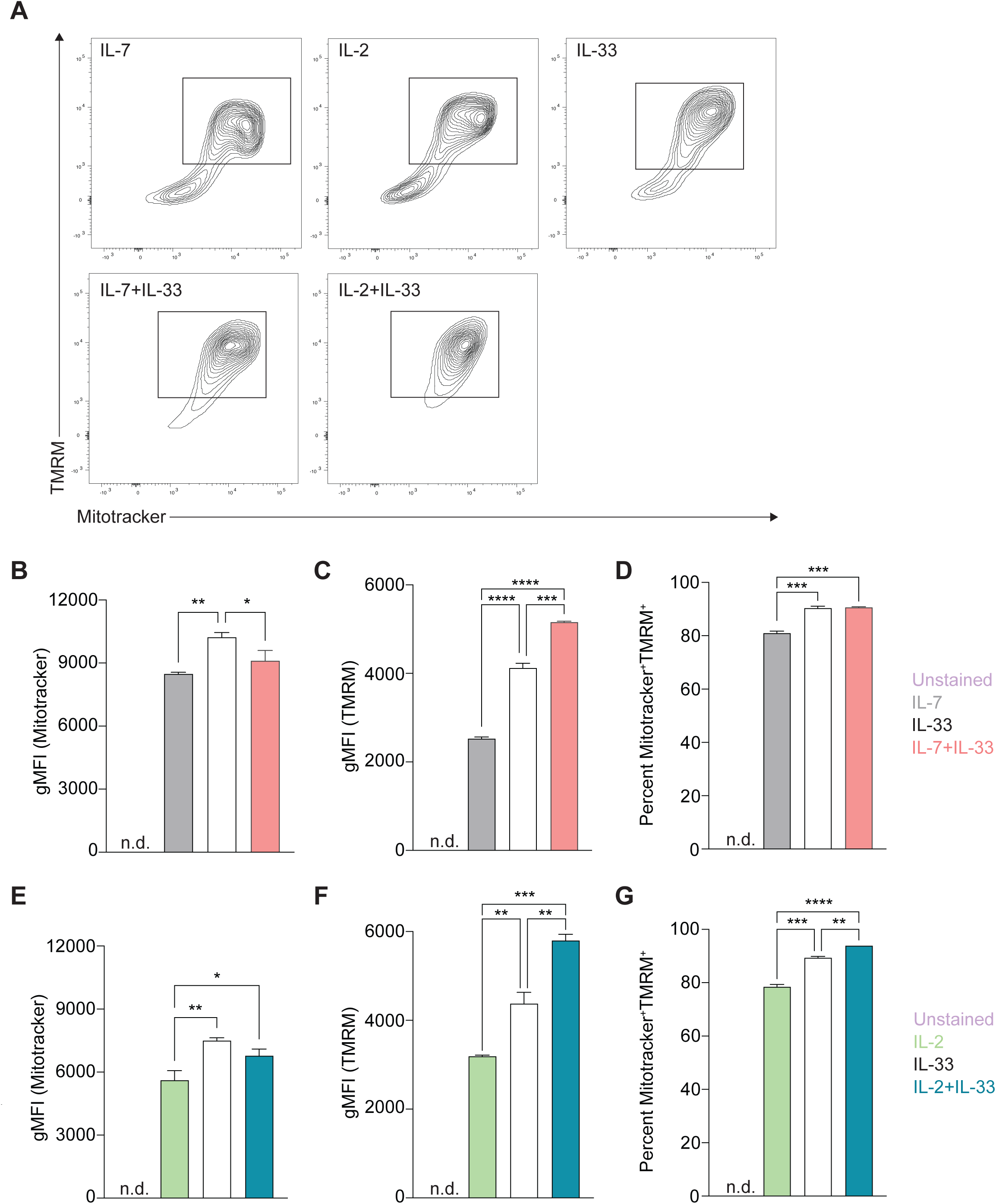
Stimulation of Group 2 innate lymphoid cells (ILC2) with activating cytokines augments mitochondrial mass and mitochondrial membrane polarization. Bone marrow-derived group 2 innate lymphoid cells (ILC2) were stimulated with either IL-7 only, IL-33 only, or a combination of IL-7 and IL-33 **(A, B-D),** or IL-2 only, IL-33 only, or a combination of IL-2 and IL-33 **(A, E-G)**. All cytokines were applied at 10 ng/mL. After 24 hours of cytokine stimulation cells were harvested and stained with Mitotracker and TMRM to quantify by flow cytometric analysis mitochondrial mass as well as mitochondrial membrane potential, respectively. ILC2 not stained with MitoTracker and TMRM were used as negative control (Unstained). Flow cytometric contour plots are depicted in **(A)** with gates set on TMRM^+^ (y-axis) and MitoTracker^+^ (x-axis) double-positive populations. From gated populations geometric Mean Fluorescence Intensities (gMFI) for Mitotracker **(B, E)** and TMRM **(C, F)**, as well as frequencies of Mitotracker^+^TMRM^+^ double-positive populations **(D, G)** were determined. Data reporting the treatment with IL-33 are the same for **(A)**, **(B-D)**, and **(E-G)**. Data are shown as average ± standard deviation (SD) and are representative of three independent experiments. Statistical analysis was performed using one-way ANOVA followed by Tukey’s multiple comparisons test (*p≤0.05, ** p≤0.01, ***p≤0.001, ****p≤0.0001).

In addition, we sought to understand how active this mitochondrial mass was by investigating their membrane polarization as an indicator of mitochondrial respiration. To further understand how cytokines known to activate ILC2 impact mitochondrial respiration and activity, bone marrow-derived ILC2 were incubated with IL-7, IL-2, IL-33 alone, or in combinations of IL-7+IL-33 or IL-2+IL-33. After 24 hours of cytokine stimulation, TMRM was added to the cells to assess mitochondrial membrane polarization, incubated for 30 minutes, followed by subsequent flow cytometric analysis (Figure 4A). Bone marrow-derived ILC2 treated with IL-7 (Figure 4C) or IL-2 alone (Figure 4F) exhibited low levels of membrane polarization, which considerably increased in the presence of IL-33. Furthermore, combined cytokine treatment of IL-7+IL-33 (Figure 4C) or IL-2+IL-33 (Figure 4F) markedly increased polarization of the mitochondrial membrane compared to either IL-7, IL-2, or IL-33 alone. These observations demonstrate that ILC2 exhibit low levels of mitochondrial polarization at steady state, actively increase their polarization upon treatment with IL-33, but especially when synergistically activated by IL-2+IL-33, or IL-7+IL-33.

Independent analysis of MitoTracker^TM^ Deep Red and TMRM yielded information about the total mitochondrial mass and the total mitochondrial membrane polarization present within each cytokine stimulation. However, the information regarding what proportion of the total mitochondrial mass was actively respirating during defined steady and activation states remained unknown. Through the implementation of gating strategies (Figure 4A), we identified mitochondria that were positive for both MitoTracker^TM^ Deep Red and TMRM to elucidate this ratio. Approximately 80% of the mitochondrial population was active in IL-7 (Figure 4D) or IL-2 alone (Figure 4G), which significantly increased to roughly 90% in the presence of IL-33. Similarly, compared to stimulations with IL-7 (Figure 4D) or IL-2 alone (Figure 4G), a marked elevation in the proportion of active mitochondria was found when ILC2 were activated by IL- 7+IL-33 (Figure 4D) or IL-2+IL-33 (Figure 4G). However, while there was a significant increase in the proportion of active mitochondria between IL-33 alone and IL-2+IL-33 (Figure 4G), this was not the case when comparing stimulations of IL-33 with IL-7+IL-33 (Figure 4D). Collectively, these observations suggest that while mitochondrial mass production plateaus during peak ILC2 activation states, the mitochondrial membrane potential and overall activity increases when synergistically activated by IL-2+IL-33.

### 3.4 Determining cell density for Seahorse Extracellular Flux Assays

We next set out to complement flow cytometry-based assays with the Agilent Seahorse XF technology to assess specific inquiries about ILC2 glucose metabolism using the Mito Stress Assay, the Glycolysis Stress Test, and the Real-Time ATP assay. These approaches generate incredibly useful data that cover a myriad of metrics; however, they are also extremely sensitive assays that are heavily influenced by cell density. As such, the number of cells present in each well play a significant role in the detection of specific measurements, as well as the consistency between replicates. If the cellular confluence within the well is too dense, this will translate to measurements that are beyond the Seahorse XF/XFe Analyzer’s detection range resulting in inaccurate readings. Likewise, cellular confluence that is too sparse will result in measurements too low to be within the detection threshold. The primary measurement that is acquired in the Mito Stress Assay is called the Oxygen Consumption Rate (OCR), and implements the use of four compounds that modulate and probe mitochondrial functions: oligomycin, carbonyl cyanide-4 (trifluoromethoxy) phenylhydrazone (FCCP), rotenone, and antimycin A. In brief, the basal OCR is measured before the injection of oligomycin, the maximum OCR value is measured after the injection of FCCP, and the lowest OCR is measured after the injection of rotenone and antimycin A. According to Agilent’s user guide *Characterizing Your Cells: Using OCR Values to Determine Optimal Seeding Density*, a cellular confluence between 50-90% and a basal OCR range of 20-160 pmol/min is recommended for optimal results. However, these recommendations are general and not specific to ILC2. To determine the optimal seeding density for all our metabolic assays, we implemented the Mito Stress Assay to evaluate which seeding density would achieve OCR values within the detectable range of the Seahorse XF/XFe Analyzer, even at their highest and lowest OCR measurements across all cytokine stimulations. We performed a titration of three different cell concentrations that were seeded and incubated for 24 hours with either IL-7, IL-2, IL-33 alone or combinations of IL-7+IL-33, as well as IL-2+IL-33. The cell seeding densities of 50,000, 75,000 or 100,000 cells per well were used in the Mito Stress Assay to obtain OCR values (Figure 5). The seeding density of 50,000 cells/well resulted in OCR values that were either not detectable or just above the detection threshold for all cytokine stimulations (Figure 5A-E, right). Similarly to 50,000 cells/well, the seeding density of 75,000 cells/well yielded OCR values that were not consistently detectable for IL-7 alone stimulations (Figure 5A, right), and were just above the detection threshold for IL-2 (Figure 5B, right), IL-33 (Figure 5C, right), IL-7+IL-33 (Figure 5D, right), and IL-2+IL-33 treatments (Figure 5E, right). The only cell seeding density that gave OCR values consistently within the detection range across all cytokine stimulations was 100,000 cells/well. Furthermore, this seeding density was also the closest to the recommended basal OCR range of 20-160 pmol/min and presented with a post-cytokine stimulation confluency of 80% (Figure 5A-E, left). As such, 100,000 cells/well was the cell seeding density chosen for all metabolic assays conducted in this paper (Figure 5A-E, left).

**Figure 5.**
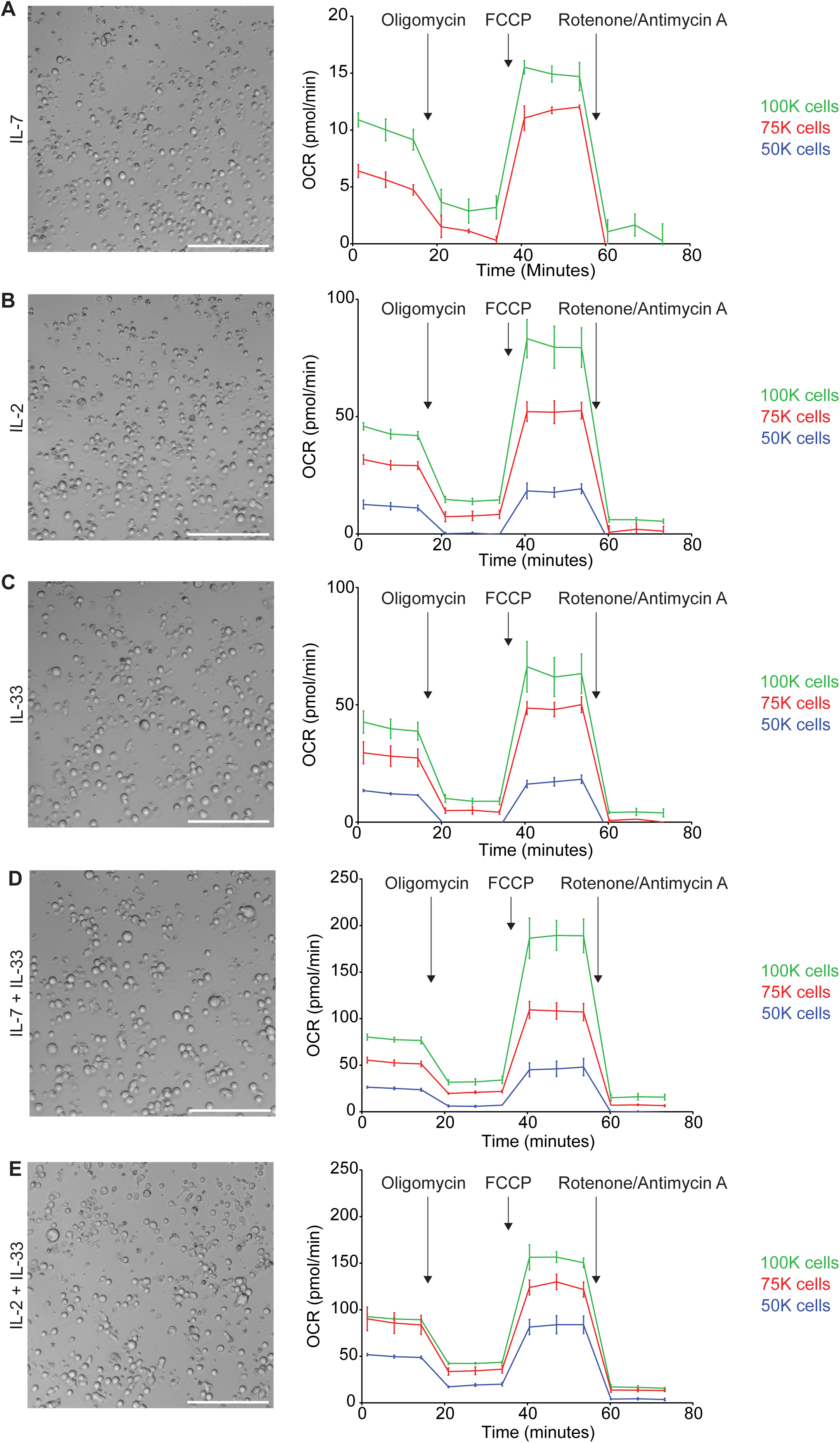
Determining cell seeding density for Seahorse metabolic assays. Bone marrow-derived group 2 innate lymphoid cells (ILC2) were seeded at three different cell densities (100.000 (100k), 75.000 (75k) or 50.000 (50k) per well) into Seahorse XFe96 microplates and stimulated with either IL-7 only **(A)**, IL-2 only **(B)**, IL-33 only **(C),** IL-7 and IL-33 **(D)**, or with IL-2 and IL-33 **(E)**. All cytokines were used at 10 ng/mL. After 24 hours of cytokine stimulation the Mito Stress Test assay was performed using the Seahorse analyzer and oxygen consumption rates (OCR) were determined. Brightfield microscopy images of seeding densities of 100.000 cells/well are shown **(A-E)**; size bars = 150mM. Data are shown as average ± standard deviation (SD) and are representative of three independent experiments.

### 3.5 Quantification of Mito Stress Assay metrics

As previously addressed in Section 3.4, the Mito Stress Assay is a metabolic assay to evaluate mitochondrial respiration and function. Mitochondrial respiration is driven by the electron transport chain (ETC); a series of five protein complexes (I, II, III, IV, V) located at the interface of the mitochondrial matrix and intermembrane space. The ETC creates an electrochemical gradient from which ATP can be produced during oxidative phosphorylation (38, 41). Through the implementation of compounds such as oligomycin, FCCP, rotenone, and antimycin A, various elements of the ETC can be manipulated to modulate mitochondrial respiration and evaluate individual metrics. The Mito Stress Assay can measure six metrics in total: basal respiration, ATP production, proton (H^+^) leak, maximal respiration, spare respiratory capacity, and nonmitochondrial respiration. Each of the metrics are calculated in reference to the OCR measurements acquired by the Seahorse XF/XFe Analyzer at specific timepoints, and in response to the compound injections that disrupt or facilitate elements of mitochondrial respiration. According to the *Seahorse XF Cell Mito Stress Test Kit User Guide* the (i) basal respiration is measured first and represents the oxygen consumption required to meet the energetic demand of the cell at baseline conditions. (ii) ATP production specifically shows the amount of ATP produced by mitochondrial respiration to meet cellular energy demands. This metric is measured when oligomycin is injected to inhibit ATP synthase (complex V), subsequently reducing the electron flow in the ETC and the subsequent OCR values. (iii) Proton leak can be an indicator of mitochondrial damage but can also be used mechanistically to regulate mitochondrial ATP production. This metric is the difference between the OCR values for basal respiration and ATP-linked respiration. (iv) The maximal respiration demonstrates the maximum rate of respiration that is possible for the cell to achieve and is measured after the injection of FCCP which mimics a physiological “energy demand.” This compound acts as an uncoupling agent that disrupts mitochondrial membrane potential by collapsing the proton gradient, enabling the ETC to operate at maximum capacity to oxidize substrates, such as glucose. (v) The spare respiratory capacity is the cell’s ability to respond to energy demands, which can be used as an indicator for cellular fitness and is calculated by subtracting the basal respiration from the maximum respiration. (vi) Finally, nonmitochondrial respiration is the cellular respiration that persists after the mitochondrial respiration has been inhibited through the injection of rotenone and antimycin A which inhibit complexes I and III, respectively.

To further understand the intricacies of mitochondrial respiration during steady and activation states, we applied the Seahorse Mito Stress Assay for sort-purified bone marrow-derived ILC2 to evaluate their proton leakage, basal respiration, maximum respiration, and spare respiratory capacity. ILC2 were incubated with IL-7, IL-2, IL-33 alone, or with combinations of IL-7+IL-33 or IL-2+IL-33. After 24 hours of cytokine stimulation, ILC2 were prepped for the Mito Stress Assay (Section 2.8) and evaluated on the Seahorse XF/XFe Analyzer (Figure 6). The levels of proton leakage in ILC2 treated with IL-7 alone were lowest of all tested cytokine treatments (Figure 6B, E). There was a marked elevation in the levels of proton leakage in the presence of IL-33 alone, as well as with the combined treatment of IL- 7+IL-33 (Figure 6B). Furthermore, the combined treatment of IL-2+IL-33 demonstrated remarkably higher proton leakage in comparison to IL-2 or IL-33 alone (Figure 6E). These observations demonstrate that ILC2 exhibit minimal signs of mitochondrial damage at steady state but were found to exhibit higher levels of proton leakage when synergistically activated by IL-7+IL-33, but especially with IL-2+IL-33.

**Figure 6.**
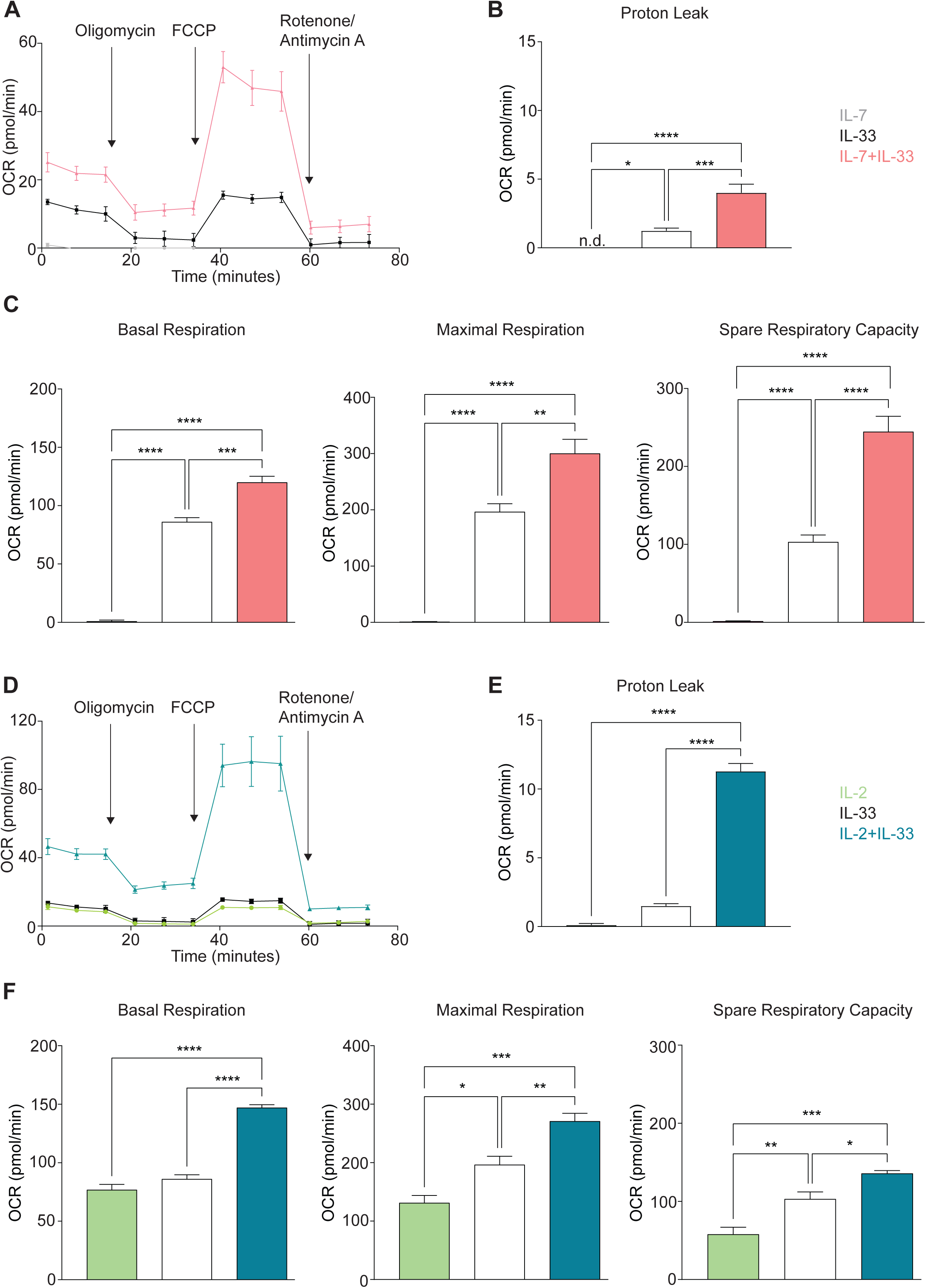
Group 2 innate lymphoid cells (ILC2) elevate oxygen consumption (OCR) rate upon stimulation with activating cytokines. Bone marrow-derived group 2 innate lymphoid cells (ILC2) were seeded into Seahorse XFe96 microplates at a cell density of 100.000 cells/well and stimulated with either IL-7 only, IL-33 only, or a combination of IL-7 and IL-33 **(A-C)**, or with IL-2 only, IL-33 only, or a combination of IL-2 and IL-33 **(D-F)**. All cytokines were used at 10 ng/mL. After 24 hours of cytokine stimulation the Mito Stress Test assay was performed using the Seahorse analyzer and oxygen consumption rates (OCR) **(A, D)**, proton leak (**B, E**), basal and maximal respiration as well as the spare respiratory capacities **(C, F)** were determined. Data reporting the treatment with IL- 33 are the same for **(A - F)**. Data are shown as average ± standard deviation (SD) and are representative of three independent experiments. Statistical analysis was performed using one-way ANOVA followed by Tukey’s multiple comparisons test (*p≤0.05, ** p≤0.01, ***p≤0.001, ****p≤0.0001).

We established in Section 3.3 that activation of ILC2 results in increased fluorescence of the dye TMRM, a dye that is preferentially acquired by mitochondria in the presence of an active membrane potential. This membrane potential is generated through the ETC during oxidative phosphorylation, and indicates mitochondrial respiration is taking place. With the Mito Stress Assay, we were then able to inhibit or facilitate specific elements of the ETC to evaluate the basal respiration, maximal respiration, and the spare respiratory capacity. These three metrics evaluated in ILC2 treated with IL-7 alone presented with minute OCR levels compared to all other cytokines (Figure 6C). There was a marked elevation in OCR levels in the presence of IL-33 alone, as well as with the combined treatment of IL-7+IL-33 (Figure 6C) for basal respiration, maximal respiration, and space respiratory capacity. In contrast, ILC2 treated with IL-2 alone already exhibited moderate OCR levels in regards to these three respiratory metrics (Figure 6F). The combined treatment of IL-2+IL-33 was significantly higher in basal respiration OCR values in comparison to treatments with IL-2, as well as IL-33 alone (Figure 6F). For both the maximal respiration and the spare respiratory capacity, the OCR values in ILC2 treated with IL-2 alone were lower in comparison to all other cytokine treatments (Figure 6F). Furthermore, there was marked elevation in the OCR values in the presence of IL-33 alone, as well as with the combined treatment of IL-2+IL-33 for the maximal respiration as well as the spare respiratory capacity. Collectively, these observations demonstrate that ILC2 are utilizing mitochondrial respiration at a minimum during steady state but were found to increase their respiration most when synergistically activated by IL-7+IL- 33, as well as IL-2+IL-33.

### 3.6 Quantification of Glycolysis Stress Test metrics

Glycolysis and oxidative phosphorylation are two major and interconnected energy producing processes in the cell. Glycolysis occurs when glucose is metabolized within the cell to generate two molecules of pyruvate, followed by a reducing reaction to form lactate as NADH and then reoxidized to make NAD^+^ (43). This reoxidization occurs in both anaerobic and aerobic glycolysis, however under anaerobic conditions this process occurs in the cytoplasm via lactate dehydrogenase (43). Under aerobic conditions the NADH is first shuttled to the mitochondria and then converted to NAD^+^ before participating in the ETC to generate ATP (43). In either case, the production of lactate in the cytoplasm results in the extrusion of H^+^ into the extracellular medium consequently raising its pH (43). In the Glycolysis Stress Test Assay, the speed at which the extracellular medium becomes acidic due this H^+^ accumulation is measured directly as the Extracellular Acidification Rate (ECAR). According to the *Seahorse XF Glycolysis Stress Test Kit User Guide,* (i) glycolysis is measured first and represents the rate of glycolysis under basal conditions. The ECAR is measured as the injected glucose is converted into pyruvate while producing water, CO_2_, NADH, H^+^, and finally ATP. (i) The glycolytic capacity is measured after the injection of oligomycin, an ATP synthase inhibitor, effectively shutting down oxidative phosphorylation and shifting energy production entirely to glycolysis. This metric exhibits the highest ECAR measurement and serves as an indication of the cells theoretical maximum rate of glycolysis. (iii) The glycolytic reserve measures the potential or ability of a cell to respond to an energy demand and is defined as the difference between the glycolytic capacity and the glycolytic basal rate. This is measured after the injection of a glucose analogue called 2-DG that inhibits glycolysis by competitively binding the first enzyme in the glycolysis pathway, causing a decrease in the ECAR measurement, which is necessary to prove that the ECAR produced in the experiment was due to glycolysis.

To investigate how cytokines known to drive ILC2 effector functions influence glycolysis during steady and activation states, we optimized the Seahorse Glycolysis Stress Test for sort-purified bone marrow-derived ILC2 to analyze basal glycolysis, glycolytic capacity, and glycolytic reserve. To this end, ILC2 were incubated with IL-7, IL-2, IL-33 alone, or with combinations of IL-7+IL-33 or IL-2+IL-33. After 24 hours of cytokine stimulation, ILC2 were prepped for the Glycolysis Stress Test (Section 2.8) and evaluated on the Seahorse XF/XFe Analyzer (Figure 7A, B). The basal glycolytic rate (Glycolysis; left) and the glycolytic capacity (centre) in ILC2 treated with IL-7 (Figure 7C) or IL-2 alone (Figure 7D) were significantly lower in comparison to all other cytokine treatments. There was a marked elevation in both metrics in the presence of IL-33 alone, as well as with the combined treatments of IL-7+IL-33 (Figure 7C) and IL-2+IL-33 (Figure 7D). In contrast, while there was no difference between IL-7 (Figure 7C) or IL-2 alone (Figure 7D) compared to IL-33 alone when measuring the glycolytic reserve values (right), the combined cytokine treatments of IL- 7+IL-33 (Figure 7C) and IL-2+IL-33 (Figure 7D) were significantly higher in comparison to all other cytokine treatments. These observations demonstrate that ILC2 are utilizing glycolysis at a minimum and exhibit a low glycolytic capacity during steady state (IL-7 or IL-2 only), but were found to increase the glycolytic rate and capacity when stimulated with IL-33, and the most when synergistically activated by IL-7+IL-33, or by IL-2+IL-33.

**Figure 7.**
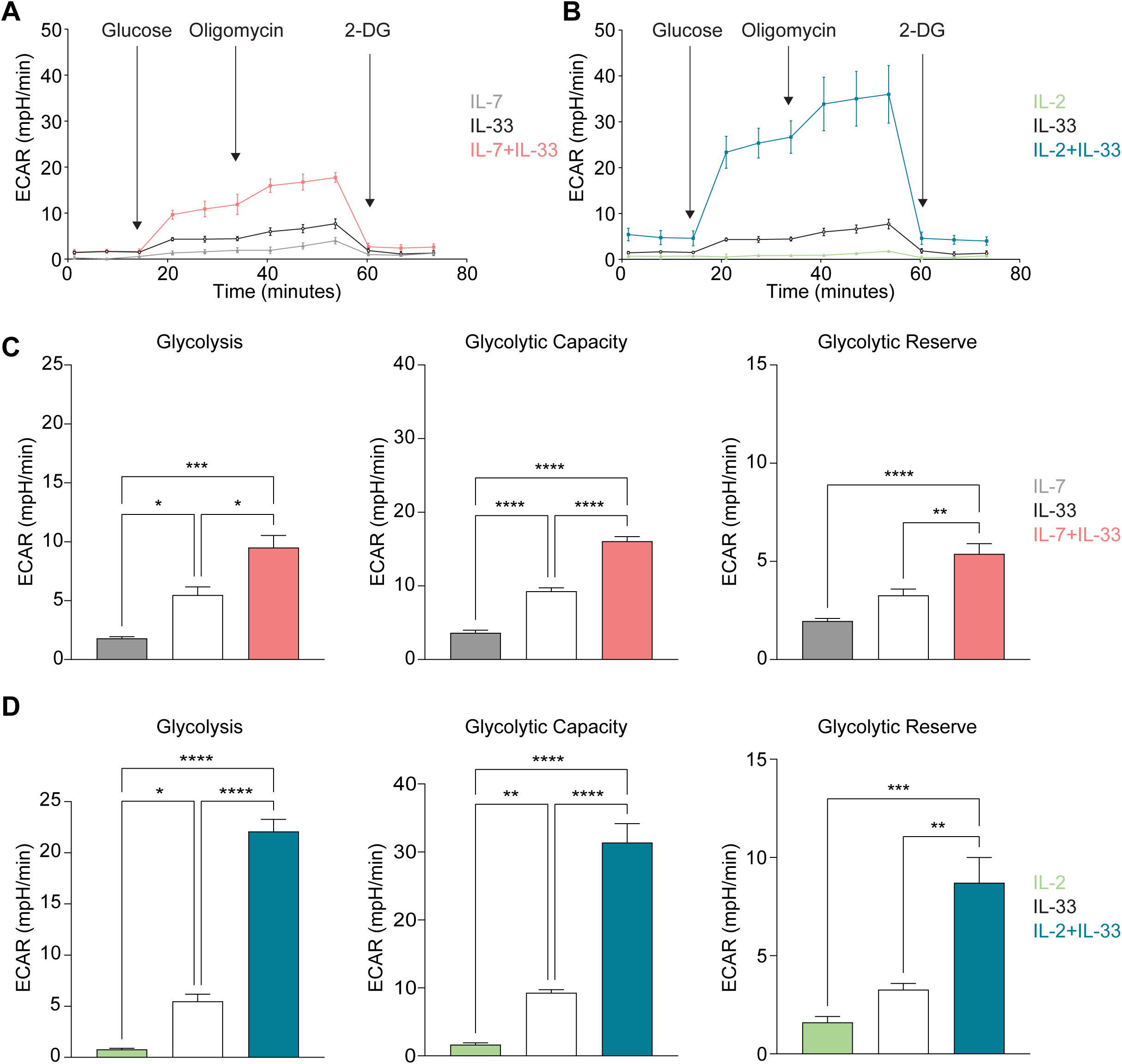
Group 2 innate lymphoid cells (ILC2) increase glycolysis upon stimulation with activating cytokines. Bone marrow-derived group 2 innate lymphoid cells (ILC2) were seeded into Seahorse XFe96 microplates at a cell density of 100.000 cells/well and stimulated with either IL-7 only, IL-33 only, or a combination of IL-7 and IL-33 **(A, C)**, or with IL-2 only, IL-33 only, or a combination of IL-2 and IL-33 **(B, D)**. All cytokines were used at 10 ng/mL. After 24 hours of cytokine stimulation the Glycolysis Stress assay was performed using the Seahorse analyzer and extracellular acidification rates (ECAR) **(A, B)**, and the levels of glycolysis, glycolytic capacity as well as the glycolytic reserve **(C, D)** were determined (2-Deoxy-D-Glucose (2-DG)). Data reporting the treatment with IL-33 are the same for **(A - D)**. Data are shown as average ± standard deviation (SD) and are representative of three independent experiments. Statistical analysis was performed using one-way ANOVA followed by Tukey’s multiple comparisons test (*p≤0.05, ** p≤0.01, ***p≤0.001, ****p≤0.0001).

### 3.7 SCENITH analysis

We next took advantage of a recently described method referred to as <u>S</u>ingle <u>C</u>ell <u>EN</u>erget<u>I</u>c metabolism by profil<u>I</u>ng <u>T</u>ranslation inhibition (SCENITH), which enables to determine global metabolic dependencies and capacities at the single-cell level. SCENITH determines the global level of translation and protein synthesis of cells, as assessed via the detection of puromycin incorporation as a primary readout, and is an especially well-suited approach to analyze the metabolism of rare cells, such as ILC2s (44). SCENITH has been shown to be reliable and comparable with other well-established techniques such as Seahorse (45). The analysis of protein synthesis in the presence of inhibitors targeting different metabolic pathways, namely 2-deoxyglucose (2-DG) for glycolysis, and oligomycin for OXPHOS, allows to assess the contribution of glycolysis vs OXPHOS for energy production of the cells. The percentage of glucose dependence quantifies how much the translation levels are dependent on glucose oxidation. As such, glucose dependence is calculated as the difference of protein synthesis levels in 2-DG treated cells compared to control (Co), divided by the difference in protein synthesis levels upon complete inhibition of ATP synthesis (2-DG + Oligomycin A (O), DGO) compared to control cells. Similarly, percentage of mitochondrial dependence quantifies how much translation is dependent on OXPHOS, which is defined as the difference in protein synthesis levels in Oligomycin A (O) treated cells compared to control relative to the decrease in protein synthesis levels upon full inhibition of ATP synthesis inhibition (DGO) also compared to control cells. Two additional parameters are calculated, ‘‘Glycolytic capacity’’ and ‘‘fatty acids and amino acids oxidation capacity’’ (FAO and AAO). Glycolytic capacity is defined as the maximum capacity to sustain protein synthesis levels when mitochondrial OXPHOS is inhibited. Conversely, FAO and AAO capacity is defined as the capacity to use fatty acids and amino acids as sources for ATP production in the mitochondria when glucose oxidation is inhibited, including glycolysis and glucose-derived acetyl-CoA by OXPHOS. While the total level of translation correlates with the global metabolic activity of the cells, the dependency parameters underline essential cellular pathways that cannot be compensated, while ‘‘capacity’’, as the inverse of dependency, reveals the maximum compensatory capacity to exploit alternative pathway/s when one is inhibited.

While global levels of translation were low when ILC2s were stimulated with IL-7 or IL-2 only, treatment with IL-33, but especially with IL-7+IL-33, and IL-2+IL-33 led to significant increase in protein synthesis (Figure 8 A, B and Figure 8E, F). Similarly, IL-33 treatment, but even more so when ILC2 were treated with IL-7+IL-33 or IL-2+IL-33 led to a reduction of glucose- as well as mitochondrial dependence, while leading to a significantly increased glycolytic, as well as FAO & AAO capacity (Figure 8 C, D and Figure 8 G, H), demonstrating a higher metabolic flexibility of ILC2 when activated by IL-33, IL-7+IL-33, and IL-2+IL-33.

**Figure 8.**
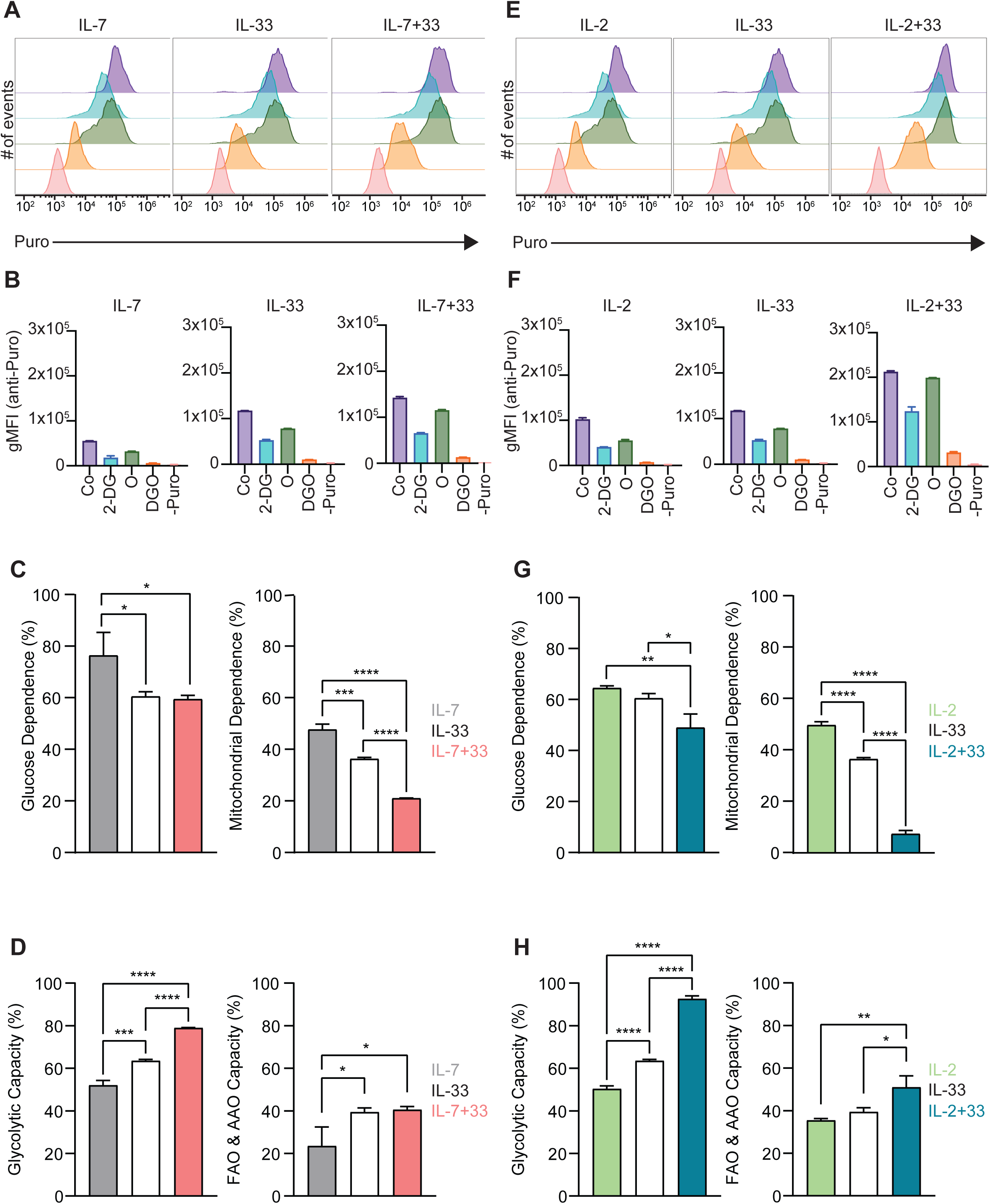
SCENITH analysis of group 2 innate lymphoid cells (ILC2). **(A-F)** SCENITH (<u>S</u>ingle <u>C</u>ell <u>EN</u>ergetIc metabolism by profil<u>I</u>ng <u>T</u>ranslation in<u>H</u>ibition) analysis was performed by stimulating bone marrow-derived group 2 innate lymphoid cells (ILC2) with either IL-7 only, IL-33 only, or a combination of IL-7 and IL-33 **(A-D),** or IL-2 only, IL-33 only, or a combination of IL-2 and IL-33 **(E-H)**. All cytokines were applied at 10 ng/mL. After 24 hours of cytokine stimulation cells were either left untreated as control (Co) or incubated with 2-Deoxy-D-Glucose (2-DG), Oligomcyin (O), or a combination of 2-DG and O (DGO) for 30 minutes. Subsequently, cells were treated for 15 minutes with puromycin followed by intracellular staining with an anti-puromycin antibody. Cells that were not treated with puromycin were used as a negative control (-Puro). ILC2 were then analyzed by flow cytometry **(A, E),** geometric mean intensities (gMFI) of the anti-puromycin staining acquired **(B, F),** and gMFI values used to determine glucose dependence and mitochondrial dependence **(C, G)**, as well as glycolytic and fatty acid oxidation (FAO) and amino acid oxidation (AAO) capacities **(D, H)**. Data reporting the treatment with IL-33 are the same for **(A-H)**. Data are shown as average ± standard deviation (SD) and are representative of three independent experiments. Statistical analysis was performed using one-way ANOVA followed by Tukey’s multiple comparisons test (*p≤0.05, ** p≤0.01, ***p≤0.001, ****p≤0.0001).

### 3.8 Characterization of ATP Production via Agilent^TM^ Seahorse

In our final analysis step, we aimed to quantify absolute ATP production levels of ILC2 using the Agilent Seahorse XF ATP Real-Time rate assay, which also measures and quantifies the rate of ATP production from the glycolytic and mitochondrial system simultaneously. As expected, stimulations with IL-7, as well as IL-2 only yielded low ATP production rates (Figure 9A, C). While treatment of ILC2 with IL-33 only significantly increased ATP production, a massive elevation of energy production was seen when cells were activated with IL-7+IL-33, or IL- 2+IL-33 (Figure 9A, C). ATP generation by ILC2 stimulated by IL-7 only largely was driven by glycolysis. In contrast, IL-2- as well as IL-33-driven energy production relied to the same extent on OXPHOS as well as glycolysis (Figure 9B, D). Compared to IL-33 stimulation only, glycolysis-mediated ATP production frequencies increased only slightly, to ∼60%, when ILC2 were treated with combinations of IL-7+IL-33 or IL-2+IL-33 (Figure 9B, D). However, the massive increase in ATP upon IL-7+IL-33 or IL-2+IL-33 stimulation, compared to IL-7, IL-2, or IL-33 only treatments, was driven by glycolysis as well as OXPHOS (Figure 9B, D).

**Figure 9.**
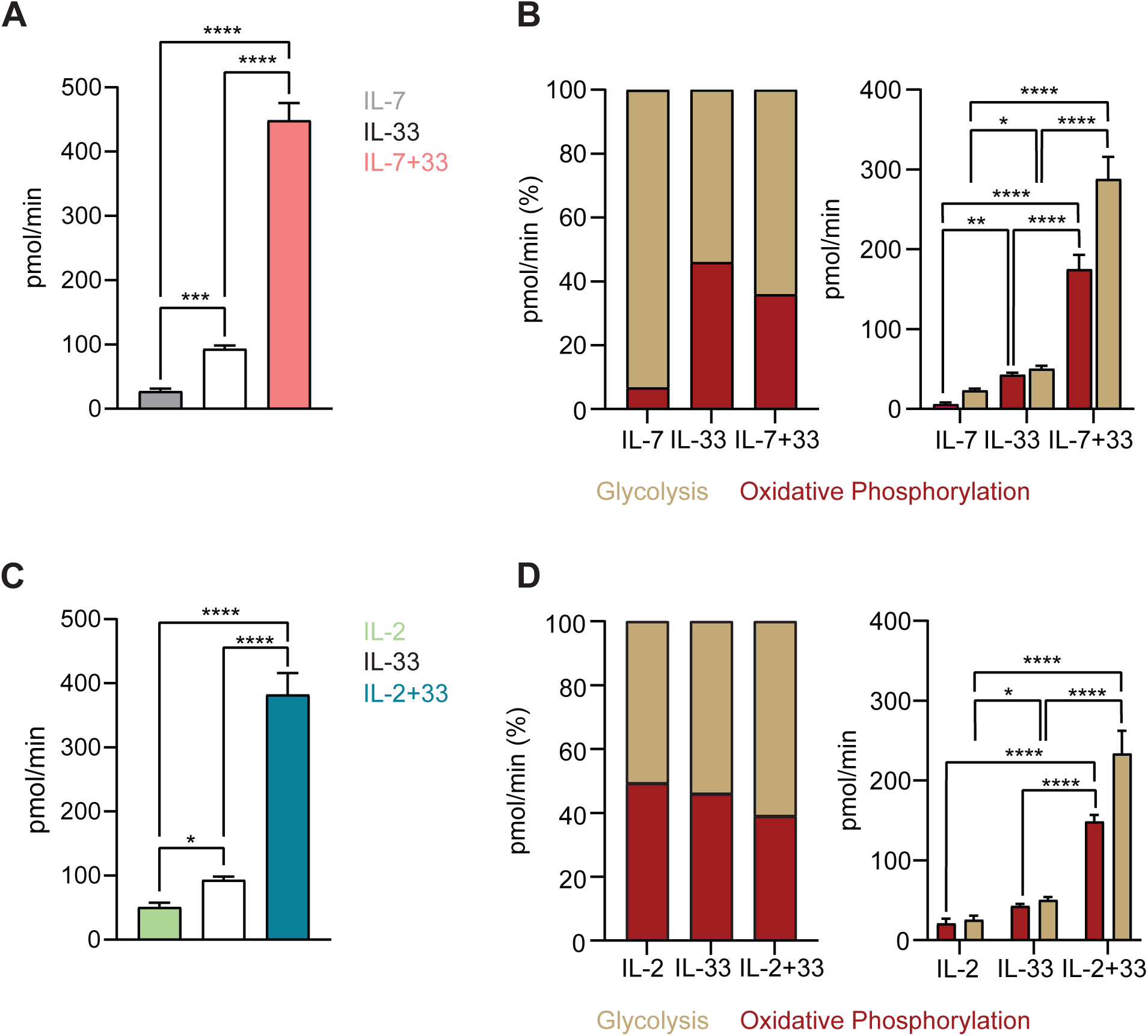
Group 2 innate lymphoid cells (ILC2) augment ATP production upon stimulation with activating cytokines. **(A-D)** To determine ATP production bone marrow-derived group 2 innate lymphoid cells (ILC2) were seeded into Seahorse XFe96 microplates at a cell density of 100.000 cells/well and stimulated with either IL-7 only, IL-33 only, or a combination of IL-7 and IL-33 **(A, B)**, or with IL-2 only, IL-33 only, or a combination of IL-2 and IL-33 **(C, D)**. All cytokines were used at 10 ng/mL. After 24 hours of cytokine stimulation the Real-Time ATP Rate assay was performed using the Seahorse analyzer. ATP production from oxidative phosphorylation as well as glycolysis were analyzed and depicted as a proportion of 100% and as absolute values. Data reporting the treatment with IL-33 are the same for **(A-D)**. Data are shown as average ± standard deviation (SD) and are representative of three independent experiments. Statistical analysis was performed using one-way ANOVA followed by Tukey’s multiple comparisons test (*p≤0.05, ** p≤0.01, ***p≤0.001, ****p≤0.0001).

## Discussion

ILC2 exert critical functions to tissue barrier integrity, driving and reinforcing immunological protection by orchestrating innate as well as adaptive immune processes. However, when deregulated, ILC2 have been shown to contribute to the pathogenesis of several chronic inflammatory barrier disorders through the release of large quantities of type 2 signature cytokines including IL-4, IL-5 and IL-13. Although recent studies started to uncover critical components of the metabolic wiring of ILC2, many aspects remain elusive. This is largely due to the inability to obtain sufficient numbers of ILC2 as well as the accessibility of rapid assays to study the intricacies of metabolic pathways. Applying our recently described protocol to expand murine bone marrow-derived ILC2 (23, 32) we detail here a framework of experimental approaches to study key immunometabolic states utilizing flow cytometry, SCENITH as well as the Seahorse platform.

We provide here an in-depth protocol of how to use ILC2 for studies applying the Seahorse platform, where treatments can be delivered to cells at specific timepoints with specific intentions to further elucidate the metabolic identities of ILC2. While the information from Seahorse assays is valuable, they require special equipment, proprietary kits and consumables, thereby limiting the number of researchers accessing this platform due to financial and accessibility restrictions. We therefore further detail flow cytometry-based assays, including SCENTITH, that can be rapidly implemented for metabolic studies of ILC2. Both, Seahorse as well as SCENITH assays, exhibit a high degree of accuracy and provided complementary insights into the metabolic wiring of ILC2. Comparatively, the high values for glycolytic and OXPHOS capacities in ILC2 treated with IL-7+IL-33 and IL-2+IL-33 found in Seahorse were corroborated by those found using SCENITH. In fact, the trend in values between single cytokine stimulations and synergistic cytokine treatments were consistent between the two platforms. In both Seahorse and SCENITH approaches, the IL-2 only conditions exhibited values that were higher than the stimulations with IL-7 alone. Although, overall SCENITH is more user-friendly and accessible to the research community, the Seahorse platform provides additional insights. As such, we would recommend, that if possible, researchers should use SCENITH to corroborate and bolster insights revealed by Seahorse.

Our here applied framework of experimental approaches combining flow cytometry assays with proprietary metabolic platforms demonstrates that the utilization of glucose is markedly elevated upon ILC2 activating cytokines. In parallel, we reveal increases in PGC1α, mitochondrial mass as well as mitochondrial membrane potential upon ILC2 activation. Our study reveals that ILC2 take up moderate levels of glucose, exhibit low levels of mitochondrial respiration, and utilize glycolysis at a minimum rate during steady state (IL-7 or IL-2 alone), but gradually increase their glucose metabolism, as well as OXPHOS upon activation with IL-33. Interestingly, while glycolysis is not heavily utilized at steady state, the proportion of ATP produced by ILC2 is almost exclusively from glycolysis in the presence of IL-7. In contrast, IL-2 treated ILC2 consistently present with a nearly 50:50 ratio of glycolysis- and OXPHOS-mediated ATP production. Overall, OXPHOS produces more ATP than glycolysis and demands a high degree of mitochondrial respiration. In fact, IL-2+IL-33 exhibited the highest values for mitochondrial respiration (TMRM) and basal respiration (Seahorse). Increased mitochondrial respiration is typically correlated with an increased demand for energy which can induce the production of ROS that damage mitochondrial integrity, consequently stimulating mitochondrial biogenesis. Concurrently, we observe that ILC2 treated with IL-2+IL-33 present the highest degree of proton leak and mitochondrial respiration. The mitochondrial mass in ILC2 treated with IL-33 was higher than that of treated with IL-7 or IL-2 alone, however, this mass decreased in the synergistic combination of these cytokines. It is possible that ILC2 treated with IL-33 increased their mitochondria production to meet the energy demand, but the increased respiration caused damage to mitochondria and the overall mass decreased as a consequence. While PGC1a levels as well as mitochondrial mass increased with IL-33 stimulation, compared to homeostatic cytokine treatments with IL-7 or IL-2 alone, synergistic activation with IL-7+IL-33 or IL-2+IL-33 did not yield their elevation above values obtained with IL-33 only. In contrast, stimulations with IL-7+IL-33 or IL-2+IL-33 markedly elevated the mitochondrial potential indicative for mitochondrial respiration. These findings were further complemented by Seahorse Real-Time ATP Rate analysis as well as SCENITH, revealing that synergistic activation of ILC2 with IL-7+IL-33 or IL-2+IL-33 led to a markedly higher ATP production and protein synthesis and elevated glycolytic and FAO & AAO capacities. Collectively this demonstrates that the increase in mitochondrial potential when synergistic activated with IL-7+IL-33 or IL-2+IL-33 translates into elevated energy production, leading to synergistic cytokine production and proliferation of ILC2 (23, 32). Interestingly, SCENITH analysis further demonstrated that ILC2 treated with IL-7+IL-33 or IL-2+IL-33 led to a reduction of glucose- as well as mitochondrial dependence, suggesting a high metabolic flexibility of ILC2 when activated by IL-33, but especially when primed by IL-7+IL-33, or IL- 2+IL-33. Collectively, these findings indicate that metabolic substrate accessibility and availability as a main driver in metabolic wiring of ILC2, which will need further experimental interrogation in future studies.

## Ethics statement

All animal experiments were performed in accordance with the guidelines and policies of the Canadian Council on Animal Care and those of McGill University.

## Author Contributions

SSK and JHF designed the intellectual framework of the study and the layout of the manuscript. SSK, AE, ARD, RCD, FR and GB performed mouse bone marrow harvests and cell culture work. SSK and AE conducted experimental work. SSK, AE and NI executed data analysis and generated Figures and Tables. SSK and RCD prepared the manuscript, which was edited by SVDR and JHF.

## Acknowledgments

The authors thank Julien Leconte and Camille Stegen from the Flow Cytometry and Cell Sorting Facility (FCCF) at the Life Sciences Complex at McGill University. The FCCF acknowledges support by the Canadian Foundation for Innovation (CFI) and by the Faculty of Medicine and Health Sciences at McGill University. The authors further thank Dr. Daina Avizonis from the McGill Metabolomics Innovation Resource (MIR) Core Facility for excellent technical support. MIR acknowledges support from the Terry Fox Foundation, CFI, Génome Québec, and a donation from the Dr. John R. Fraser and Mrs. Clara M. Fraser Memorial Trust.

## Publisher’s note and Copyright

All claims expressed in this article are solely those of the authors and do not necessarily represent those of their affiliated organizations, or those of the publisher, the editors and the reviewers. Any product that may be evaluated in this article, or claim that may be made by its manufacturer, is not guaranteed or endorsed by the publisher.

This is an open-access article distributed under the terms of the Creative Commons Attribution License (CC BY). The use, distribution or reproduction in other forums is permitted, provided the original author(s) and the copyright owner(s) are credited and that the original publication in this journal is cited, in accordance with accepted academic practice. No use, distribution or reproduction is permitted which does not comply with these terms.

---

**Supplemental Figure S1.** Gating strategies for the isolation of murine bone marrow-derived group 2 innate lymphoid cell (ILC2). Debris and doublets were excluded and murine bone marrow-derived ILC2 precursors were defined and isolated by flow cytometric sorting as lineage-negative, c-kit^-^Sca-1^+^CD25^+^ cells.

