## Supplementary figures and images for "Immunometabolic analysis of primary murine Group 2 Innate Lymphoid Cells: a robust step-by-step approach"

### Supplemental Figure 1

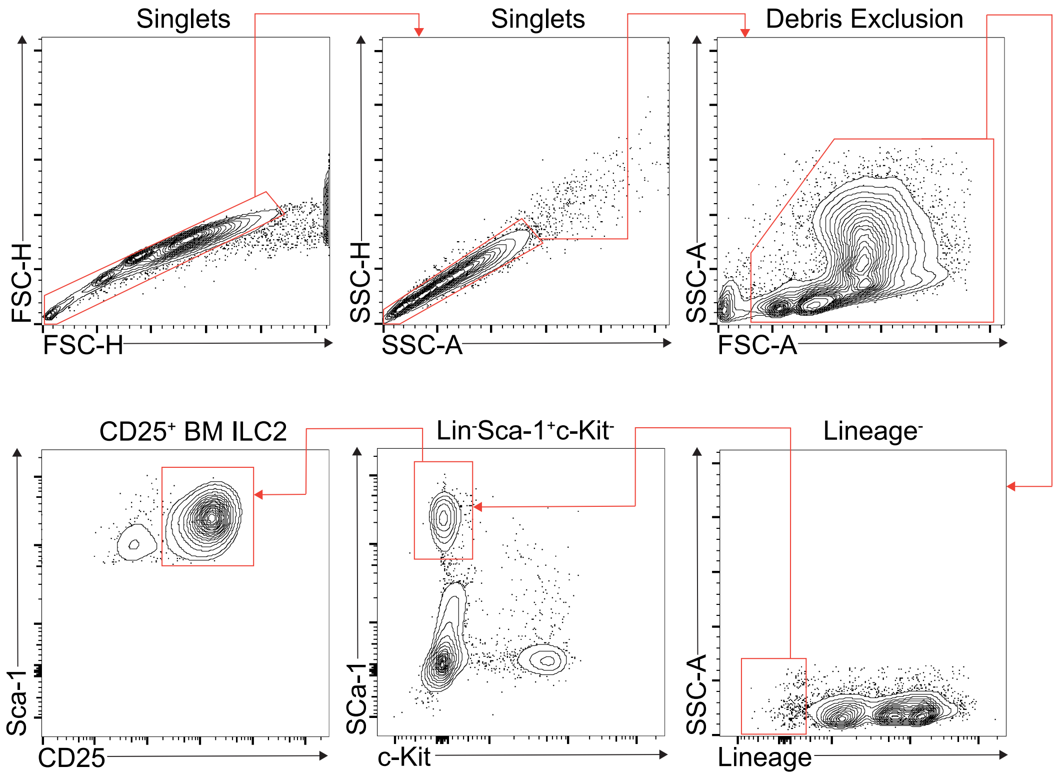


Supplementary Figure 1
